# Dual plasmepsin IX and X inhibitors are refractory to development of resistance

**DOI:** 10.64898/2026.03.09.710546

**Authors:** Paola Favuzza, Madeline G. Dans, Wenyin Su, Jennifer K. Thompson, Anthony N. Hodder, Anna Ngo, Jocelyn Sietsma Penington, Danushka S. Marapana, Tony Papenfuss, Manuel de Lera Ruiz, Rachael Coyle, Marcus C. S. Lee, John A. McCauley, Kym Lowes, David B. Olsen, Brad E. Sleebs, Alan F. Cowman

## Abstract

Artemisinin-based combination therapies (ACTs) remain the cornerstone of malaria treatment, but emerging resistance threatens their efficacy. The potential for the development of drug resistance against plasmepsin X (PMX)-selective inhibitors and dual plasmepsin IX/X (PMIX/X) inhibitors was investigated in *Plasmodium falciparum*. A series of PMX-selective (WM4, WM76, WM92) and PMIX/X dual inhibitors (WM382, WM09, WM42) were characterised for potency against parasite growth and enzyme inhibition. *In vitro* selection experiments showed that all compounds had a high barrier to resistance, although parasites with reduced sensitivity to PMX-selective inhibitors could still be selected. Resistance mechanisms involved *pmx* gene amplification and point mutations (D245N, S315P, S359P, I363L) that alter inhibitor binding. Recombinant expression and Michaelis–Menten kinetics demonstrated that these mutations impair drug binding whilst preserving PMX catalytic function. Reverse genetics confirmed that introducing these mutations into the *pmx* gene resulted in decreased potency of the inhibitors. In this study, resistance to the PMIX/X dual inhibitors evaluated here could not be selected, despite prolonged selection pressure. Antimalarial Resistome Barcoding (AReBar) assays confirmed the absence of pre-existing resistance to either inhibitor class. Critically, PMIX/X dual inhibitors maintained efficacy against parasites with decreased sensitivity to PMX-selective compounds. These findings demonstrate that dual PMIX/X inhibitors present a substantially higher barrier to resistance than PMX-selective inhibitors, informing antimalarial drug development strategies and highlighting dual-target inhibition as a promising approach to mitigate resistance risks.

## Introduction

The global fight against malaria has faced significant challenges in recent years, despite notable progress in the early 21st century. From 2000 to 2019, annual malaria deaths declined substantially from 864,000 to 576,000 [1]. However, the COVID-19 pandemic reversed this positive trajectory, with mortality increasing to 631,000 in 2020 and remaining elevated at 610,000 deaths in 2024 - 43,000 more than pre-pandemic levels. Similarly, global malaria cases rose from 233 million in 2019 to approximately 282 million in 2024, highlighting the pandemic’s devastating impact on malaria control efforts [1].

Artemisinin-based combination therapies (ACTs) continue to be the cornerstone of malaria treatment, particularly for *Plasmodium falciparum* infections. The scale of ACT deployment is remarkable, with approximately 4.7 billion treatment courses distributed globally between 2010 and 2024, including 260 million in 2024 alone [1]. However, this widespread use has been accompanied by an emerging issue: the development of complex, multifaceted drug resistance mechanisms. Initially identified in Southeast Asia nearly two decades ago, artemisinin-resistant *P. falciparum* has now been documented in South America (Guyana), Papua New Guinea, and most critically, has emerged independently in East Africa. Countries including Rwanda, Uganda, South Sudan, Tanzania, Ethiopia, Eritrea, and eastern Democratic Republic of Congo (DRC) have reported established artemisinin resistance (reviewed in [2, 3]). Artemisinin resistance is complex but primarily mediated through mutations in PfKelch13 that affects the protein’s regulatory function in cell stress responses [4, 5] that act together with other mutations in proteins involved in the endocytic pathway (reviewed in [6]). The partner drugs of artemisinin in ACTs are also succumbing to resistance, with increased gene copy numbers of *P. falciparum plasmepsin 2* and *3* correlating with piperaquine resistance in Southeast Asia [7-9].

The discovery and development of novel antimalarial drugs have become increasingly important due to the rise of *P. falciparum* resistance to current frontline treatments. Several promising new drug targets are being explored, including *Plasmodium* proteasome inhibitors, which have shown efficacy against both blood and liver stages of the parasite [10]. Other approaches target the parasite’s phosphatidylinositol 4-kinase (PI4K), as exemplified by MMV390048 which entered clinical trials [11], and the enzyme dihydroorotate dehydrogenase (PfDHODH) with compounds like DSM265 showing potential against drug-resistant strains [12]. Another promising candidate is Ganaplacide, that has recently been developed in combination with lumefantrine as GanLum, showing potential as a novel antimalarial treatment [13]. The focus on novel targets is essential because of the ability of *P. falciparum* to evolve new resistance mechanisms. This has spurred increased investment in drug candidates with entirely new mechanisms of action that would potentially circumvent existing resistance pathways [14].

Aspartic proteases in *Plasmodium* species, particularly plasmepsin IX (PMIX) and plasmepsin X (PMX), have emerged as therapeutic targets against malaria [15-19]. These proteases play essential roles in the parasite life cycle, with PMIX being required for merozoite egress through involvement in subtilisin 1 maturation, and PMX functioning as a maturase for SERA5 and SERA6 during schizont rupture [15-17]. Additionally, PMX processes a number of *P. falciparum* proteins that play important roles in merozoite invasion including the essential ligand PfRh5 [20, 21]. The indispensable nature of these enzymes for parasite survival, combined with distinct structural features compared to human aspartic proteases, has led to development of new antimalarial compounds, notably with the discovery of WM382, a potent dual inhibitor that simultaneously targets PMIX and PMX [17, 18].

Recent structural and biochemical studies have revealed that PMIX and PMX possess distinctive substrate specificity pockets that differ significantly from their human homologs [22]. This structural divergence has been leveraged in drug discovery efforts, leading to the evolution of WM382 to MK-7602, a promising clinical candidate maintaining dual inhibition of PMIX and PMX. The discovery of MK-7602 demonstrates the feasibility of targeting these essential proteases with antimalarials that could lead to new therapeutic options with both curative and transmission-blocking properties. Both WM382 and MK-7602 target the liver, blood and transmission stages of *P. falciparum* and they have a high barrier for the selection of resistance due to their dual targeting of PMIX and PMX [17, 19]. The pharmacokinetic profile and tolerability of MK-7602 have been evaluated in two phase 1 studies and support continued clinical development [23]. Additionally, the antimalarial activity of clinical candidate MK-7602 has been tested in a phase 1b Controlled Human Malaria Infection (CHMI) human clinical trial with encouraging results (NCT06294912).

The analysis of resistance mechanisms for new antimalarial clinical candidates involves a multi-layered approach combining *in vitro* evolution studies, genomic analysis, and clinical surveillance. This comprehensive resistance analysis has become important for early identification of resistance mechanisms to assist in optimizing dosing strategies to minimize resistance development. In *P. falciparum* this approach has successfully identified mutations in PfCARL as a resistance mechanism for the imidazolopiperazine compound Ganaplacide, and mutations in PfATP4 conferring resistance to several new antimalarial candidates including Cipargamin [24, 25].

In this study, the possibility of selecting resistance to PMX selective inhibitors and PMIX/X dual inhibitors by *P. falciparum* was investigated. This included the PMX selective inhibitors WM4, WM76, WM92 that were compared to the dual PMIX/X inhibitors WM382, WM09 and WM42. Our findings demonstrated a high barrier against the selection of resistance in *P. falciparum*; however, it was possible to select decreased sensitivity for the PMX selective inhibitors. The molecular mechanisms of resistance involved amplification of the *pmx* gene and mutations within the protease that alter the ability of the inhibitors to bind and inhibit protease activity were identified. This has contributed to our understanding of potential resistance mechanisms to these promising new antimalarial drug candidates and informs strategies for their optimal development and deployment.

## Results

### Characterization of selective and dual inhibitors targeting P. falciparum plasmepsin IX and X

A high-throughput screen of aspartic protease inhibitors identified iminopyrimidinone compounds that blocked *P. falciparum* growth, including the hit compound WM4 (Fig 1A) [17, 18]. This discovery prompted a medicinal chemistry campaign to develop structural analogs of WM4, ultimately leading to the tool compound WM382 and clinical development compound MK-7602, both potent dual inhibitors of PMIX and PMX [19]. The program involved the design and synthesis of approximately 5,000 compounds, including WM382, WM76, WM09, WM92, and WM42 (Fig 1A) [17, 18].

**Figure 1.**
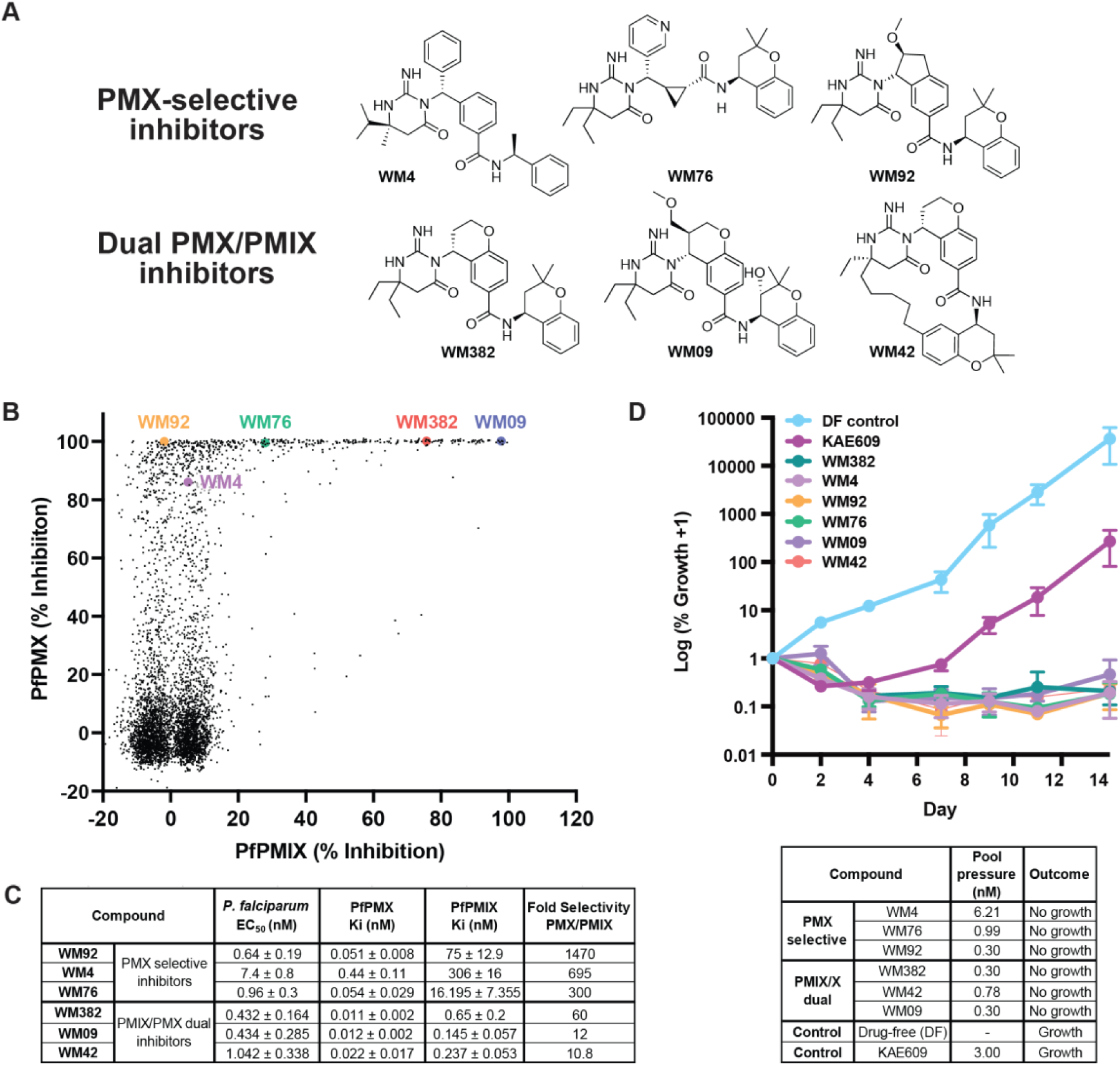
PMX selective and dual PMIX/X inhibitors. **A.** Chemical structures of PMX-selective and PMX/IX dual inhibitors. **B.** A library of 4505 aspartic protease inhibitors (10 nM) was screened against recombinant *P. falciparum* PMX (0.2 nM) and PMIX (5 nM) and plotted as percent inhibition of enzyme activity. **C.** EC_50_ for *P. falciparum* and K_i_ for PfPMX and PfPMIX (Mean ± SD, three independent experiments). **D.** PMX-selective and PMIX/X dual inhibitors profiled in AReBar assay at concentrations equivalent to 3x EC_50_, alongside drug-free treatment (DF, light blue) and KAE609 (magenta) as controls. DF control, KAE609, WM4 and WM382 were previously described [19]. Concentrations of compounds used in the experiment (pool pressure) are reported in the lower table.

All synthesized compounds were systematically evaluated for their inhibitory activity against recombinant *P. falciparum* aspartic proteases PMIX and PMX (Fig 1B) [17]. This extensive screening effort revealed three distinct groups of compounds: those showing little or no inhibitory activity against either PMIX or PMX, PMX-specific inhibitors including WM4, WM76, and WM92, and PMIX/X dual inhibitors WM382 and WM09 (Fig 1B, C). Notably, no PMIX-selective compounds were identified during this process (Fig 1B), highlighting the unique properties of these aspartic proteases and challenges in discovering selective inhibitors.

PMX-selective and dual PMIX/X inhibitors demonstrated potent inhibition of *P. falciparum* asexual blood stage growth (Fig 1C), consistent with the essential nature of both PMIX and PMX in the parasitic lifecycle [15-17]. The optimized PMX-selective inhibitors WM92 and WM76 showed significantly increased potency against *P. falciparum* blood stage parasites compared to the hit compound WM4, with EC_50_ values of 0.64 nM and 0.96 nM, respectively (Fig 1C). Similarly, the dual PMIX/X inhibitors displayed comparable potency, with WM09 and WM42 exhibiting EC_50_ values of 0.43 nM and 1.04 nM, respectively. These optimized compounds demonstrated substantially improved potency over the hit compound WM4 (EC_50_ 7.4 nM) (Fig 1C) and were comparable to the tool compound WM382 (0.43 nM) and the clinical candidate MK-7602 [17, 19].

To further characterize the compounds, we determined their inhibitory binding constants (K_i_) for PMIX and PMX, providing insight into their potency and selectivity profiles (Fig 1C). The hit compound WM4 selectively inhibits PMX with a K_i_ of 0.44 nM, demonstrating a 695-fold selectivity over PMIX [17, 22]. The optimized PMX-selective inhibitors WM76 and WM92 showed even greater potency and selectivity, with PMX K_i_ values of 0.054 nM and 0.051 nM, respectively, and selectivity ratios against PMIX of 300-fold and 1,470-fold (Fig 1C). In contrast, the dual inhibitors WM382, WM09, and WM42 exhibited potent inhibition of both proteases, albeit with varying degrees of selectivity. The lead compound WM382 showed a 60-fold selectivity between PMX and PMIX, while WM09 and WM42 demonstrated K_i_ values for PMX of 0.01 nM and 0.02 nM, with selectivity ratios of 12-fold and 10.8-fold, respectively. These results highlight the development of both highly selective PMX inhibitors and potent dual PMIX/X inhibitors, providing a set of tools to further investigate the function of these enzymes during different stages of parasite replication. Moreover, these compounds provide valuable tools to examine the differential propensity for resistance development in *P. falciparum* between PMX-specific and PMIX/X dual inhibitors.

### PMIX and PMX inhibitors demonstrate no cross-resistance with known mechanisms of drug resistance

To investigate potential cross-resistance between PMX-selective and PMIX/X dual inhibitors and known antimalarial resistance mechanisms, the Antimalarial Resistome Barcoding (AReBar) library of resistant *P. falciparum* parasites was employed (Fig 1D) [26]. This comprehensive approach simultaneously assesses the efficacy of the compounds against a population of 52 *P. falciparum* parasite lines possessing genetically engineered to express resistance mechanisms to a broad range of compounds, accounting for most known resistance mechanisms (S1 Table). EC_50_ values were determined for inhibition of *P. falciparum* Dd2 and 3D7 growth to determine the pool pressure (Fig 1D) to be applied during the experiment.

Over the course of the 14-day AReBar assay, the drug-free control population expanded as expected (Fig. 1D). In contrast, parasites exposed separately to 3x EC_50_ of the PMX-selective inhibitors (WM4, WM76, and WM92) and PMIX/X dual inhibitors (WM382, WM09, and WM42) failed to grow, with no viable parasites detected at the end of the selection period. This contrasted with the control compound Cipargamin (also known as KAE609), a PfATP4 inhibitor, which allowed for the expansion of *P. falciparum* lines encoding resistance mechanisms specific to this drug [17, 27]. The Cipargamin-resistant parasites were readily detectable by day 14, serving as a positive control for the assay’s ability to identify resistant populations (Fig. 1D). The lack of growth in the presence of our PMIX and PMX inhibitors demonstrates that known mechanisms of resistance in *P. falciparum* do not confer any protection against these novel compounds. This outcome, while not entirely unexpected given the unique mechanisms of action of PMIX and PMX inhibitors, provides evidence for their potential to overcome existing in-the-field clinical resistance [17].

### PMX-selective and PMIX/X dual inhibitors have a high barrier to drug resistance development in P. falciparum

PMX-selective and PMIX/X dual inhibitors provided valuable tools to compare the risk of resistance for compounds targeting single versus dual targets in *P. falciparum*. The established method of exposing *P. falciparum* blood stage parasites to drug pressure using a slow, incremental increase in the concentration of each tested compound was employed [28, 29]. Drug selection with the hit compound WM4 for 6 months yielded a parasite line with an EC_50_ of 66.6 nM corresponding to a 6-fold shift (Fig 2A). Using the same approach, parasites with decreased sensitivity to the selective PMX inhibitors WM76 and WM92 were generated (Fig 2B, C) over 15 and 3 months, achieving an approximate 28-and 42-fold shift in EC_50_, respectively (WM76-selected EC_50_ 23.5 nM and WM92-selected EC_50_ 34.8 nM) (Fig 2B, C). In stark contrast, dual PMIX/X inhibitors WM09 and WM42 demonstrated a resistance profile similar to WM382, where neither decreased sensitivity nor resistance could be generated even after prolonged selection (Table 1) [17].

**Figure 2.**
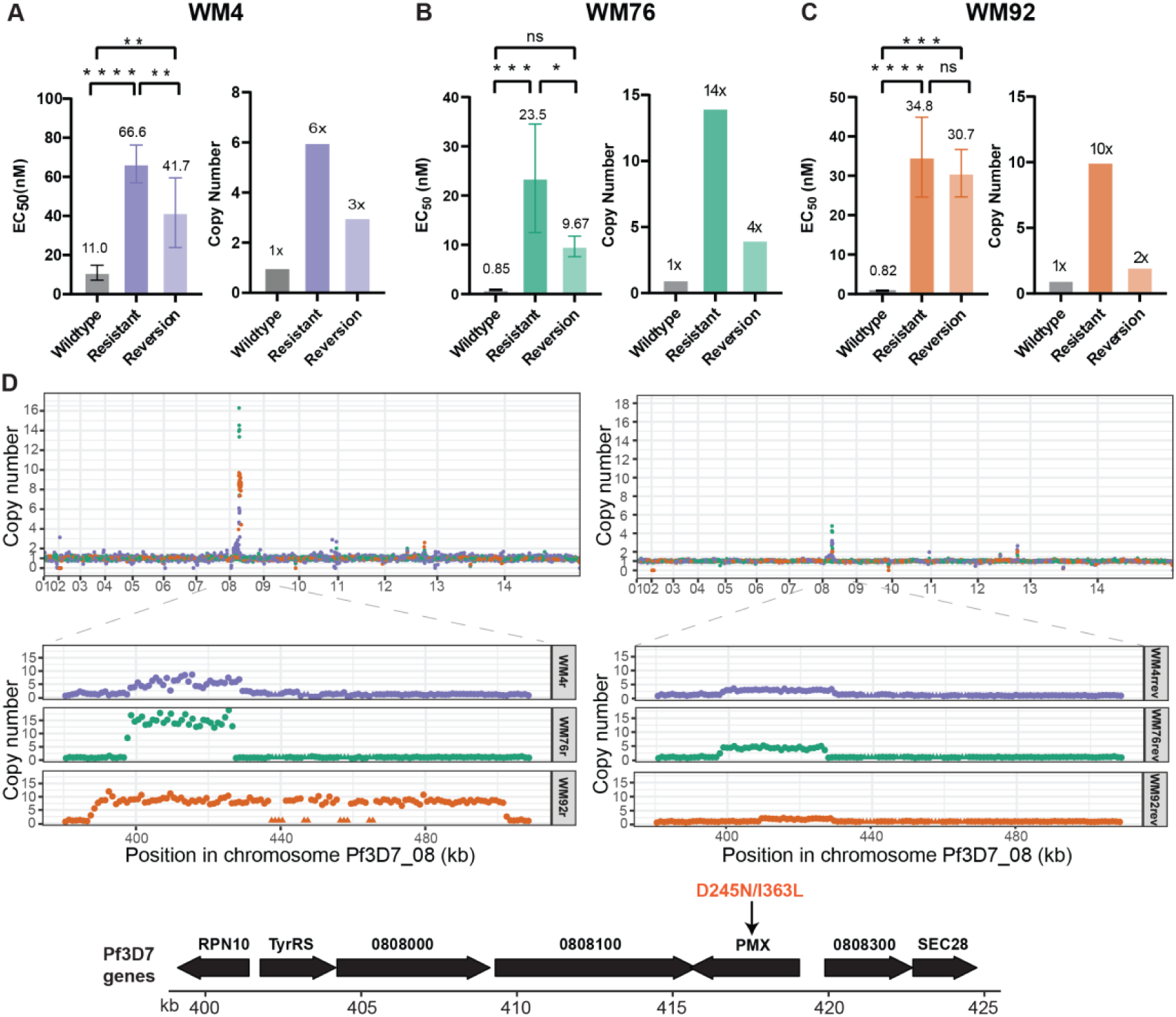
Resistance generated against PMX-selective inhibitors. **A.** Left panel shows the EC_50_ values (nM) derived from growth inhibition assay for wildtype, WM4-resistant and WM4-reversion *P. falciparum* lines. The right panel shows the *pmx* gene copy number for these parasite lines. **B.** Left panel shows the EC_50_ values (nM) derived from growth inhibition assays for wildtype, WM76-resistant and WM76-reversion *P. falciparum* lines. The right panel shows the *pmx* gene copy number for these parasite lines. **C.** Left panel shows the EC_50_ values (nM) derived from growth inhibition assays for wildtype, WM92-resistant and WM92-reversion *P. falciparum* lines. The right panel shows the *pmx* gene copy number for these parasite lines. For A, B, and C the error bars represent the SD of >3 independent experiments. Statistical analysis conducted by Ordinary One-way ANOVA with multiple comparisons in GraphPad Prism. * p<0.05, ** p<0.01, *** p< 0.001, **** p<0.0001. **D.** Genome of *P. falciparum* WM4-, WM76- and WM92-resistant (left) and reversion (right) lines showing copy number (top) and detail of amplification of chromosome 8 (bottom). Triangles indicate regions where copy number was not determined. Gene numbers identified as amplified in this region can be found at PlasmoDB: https://plasmodb.org/plasmo/app/. WM4 (purple), WM76 (green), WM92 (orange). The D245N/I363L SNPs identified in the *pmx* gene are highlighted.

**Table 1.**
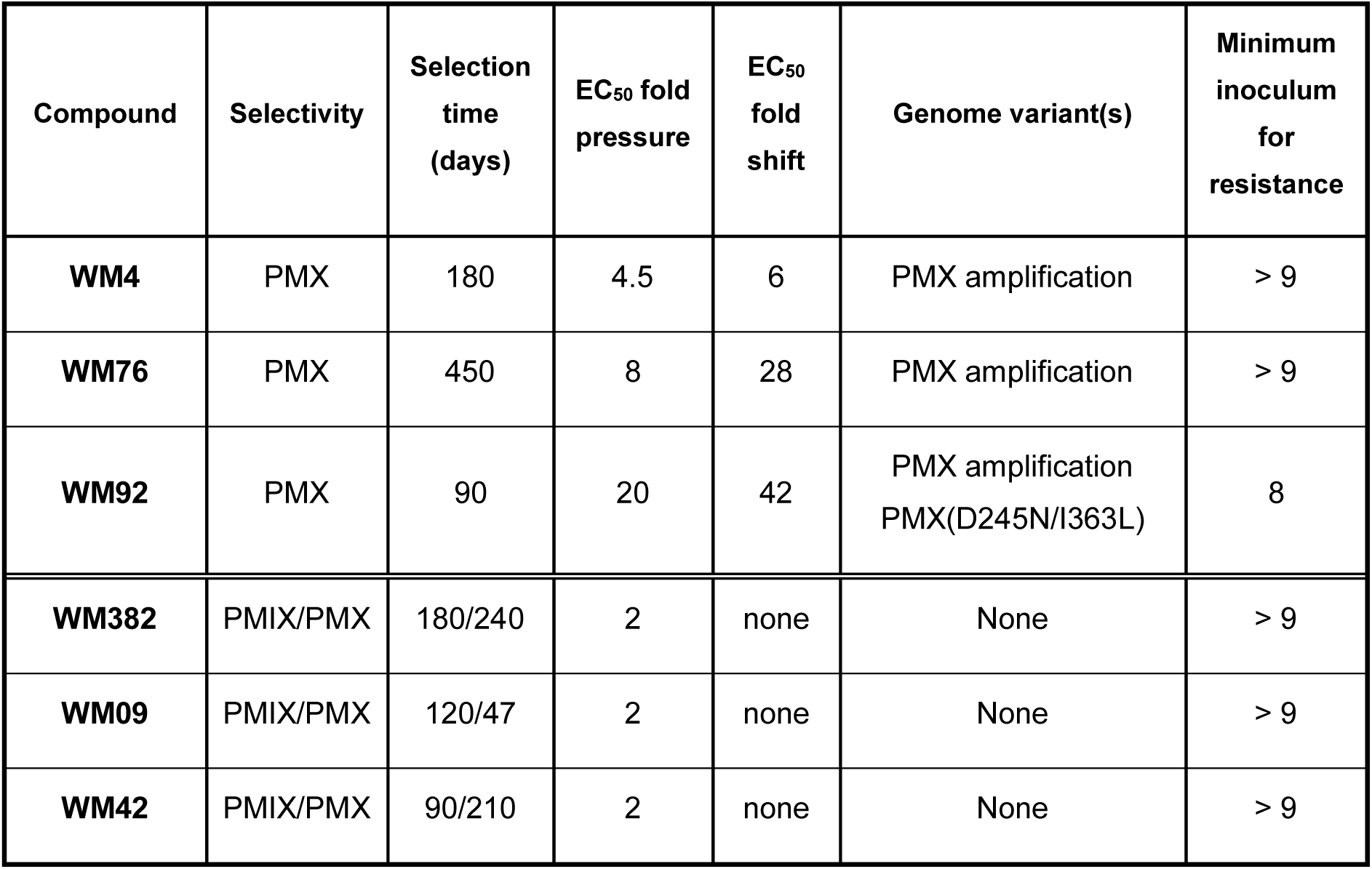
Resistance profile characterization with PMX selective and PMIX/X dual inhibitors.

| Compound | Selectivity | Selection time (days) | EC<sub>50</sub> fold pressure | EC<sub>50</sub> fold shift | Genome variant(s) | Minimum inoculum for resistance |
| --- | --- | --- | --- | --- | --- | --- |
| **WM4** | PMX | 180 | 4.5 | 6 | PMX amplification | > 9 |
| **WM76** | PMX | 450 | 8 | 28 | PMX amplification | > 9 |
| **WM92** | PMX | 90 | 20 | 42 | PMX amplification<br>PMX(D245N/I363L) | 8 |
| **WM382** | PMIX/PMX | 180/240 | 2 | none | None | > 9 |
| **WM09** | PMIX/PMX | 120/47 | 2 | none | None | > 9 |
| **WM42** | PMIX/PMX | 90/210 | 2 | none | None | > 9 |

To elucidate the mechanisms underlying decreased sensitivity to the PMX selective inhibitors, whole genome sequencing was performed on WM4-, WM76-, and WM92-selected *P. falciparum* lines. This analysis revealed increased copy number of the genomic region surrounding the *pmx* gene on chromosome 8 in all three lines (Fig 2A-D). The WM4-selected line showed up to 6 copies of a *pmx*-containing amplicon, consistent with our previous findings [17]. The WM76-selected line exhibited a 14-fold increase in a 15 kb amplicon, with no associated SNPs identified. The WM92-selected line displayed approximately 10 copies of a 28 kb amplicon that included the *pmx* gene. Two SNPs in the *pmx* gene were also identified, resulting in D245N and I363L amino acid changes within the PMX protease (S1 Fig). These findings demonstrated that slow incremental drug pressure could select for decreased sensitivity to PMX-selective compounds through both gene amplification and, in the case of WM92, amino acid changes in the PMX protease. Notably, the inability to select for decreased sensitivity or resistance to dual PMIX/X inhibitors WM382, WM09 and WM42 suggests a significantly higher barrier to resistance for these compounds compared to PMX selective inhibitors.

To assess potential fitness costs associated with *pmx* gene amplification and mutations, WM4-, WM76-, and WM92-selected lines were cultured in drug-free media for 6 months. Subsequent drug sensitivity analysis revealed significantly reduced EC_50_ values for WM4 and WM76 against the respective compounds used for selection (Fig 2A, B). The WM4-reversion line showed an EC_50_ of 41.7 nM down from 66.6 nM (Fig 2A), while the WM76-reversion line exhibited an EC_50_ of 9.67 nM down from 23.5 nM (Fig 2B). Intriguingly, the WM92-reversion line maintained essentially the same EC_50_ (30.7 nM) as the resistant line (34.8 nM) (Fig 2C).

Genome sequencing of each reversion line revealed reductions in *pmx* gene copy number. The WM4-reversion line had a 2-fold reduction (from 6 to 3 *pmx* copies), and the WM76-reversion line demonstrated a 3.5-fold decrease (from 14 to 4 copies of the *pmx* gene) (Fig 2A, B, D). Surprisingly, although the WM92-reversion line did not show a reduced EC_50_ to WM92 compared with the WM92-resistant line (Fig 2C), the reversion parasite line exhibited a 5-fold decrease in *pmx* gene copy number (from 10 to 2 copies of the *pmx* gene) (Fig 2C, D). The presence of both mutations in both WM92-resistant and WM92-reversion parasite lines was confirmed by PCR using *pmx*-specific primers (S1 Fig, left panel) and by Sanger sequencing of the amplified products (S1 Fig, right panels). These findings suggested that the D245N/ I363L amino acid change plays a key role in conferring and maintaining decreased sensitivity to WM92, even after a reduction in gene copy number.

During these resistance studies, a ‘long-term selected’ parasite line was also generated with wildtype parasites exposed to increasing amounts of WM4 up to 250 nM for over two years. This resulted in an EC_50_ shift of 156-fold (653 nM compared to 4.20 nM in 3D7) (S2A Fig). Whole genome sequencing of this long-term selected line revealed a 14x copy number increase surrounding the *pmx* gene, with an additional SNV identified as S315P in PMX (S2B, C Fig).

Together, these results indicate that although the selection of resistance to PMX-selective inhibitors was possible, it is difficult to achieve and often unstable. In contrast, resistance to dual PMIX/X inhibitors appears even more challenging to select, reflecting a more robust resistance profile. This highlights potential advantages of multi-target inhibition strategies in antimalarial drug development and underscores the complex interplay between gene amplification and point mutations in conferring drug resistance in *P. falciparum*.

### Determination of the minimum inoculum for resistance

To further evaluate the propensity for the development of resistance, minimum inoculum for resistance (MIR) studies were conducted for WM4, WM76, WM92, and WM382, WM09 and WM42, using the *P. falciparum* Dd2 line. This approach involved exposing a range of *inocula* of parasites to drug pressure for 60 days and determining the smallest parasite *inoculum* from which resistant mutants can be selected [30, 31]. The dual PMIX/X inhibitors WM382, WM09 and WM42 demonstrated exceptional barrier to the development of decreased sensitivity. Even at the highest *inoculum* of 10^9^ parasites, no recrudescence was observed over the 60-day selection period, indicating an MIR >9 (Fig 3A, Table 1). This result underscores the high barrier to resistance for these dual-target inhibitors. Interestingly, the PMX-selective compounds WM4 and WM76 also exhibited a high barrier to resistance in this assay, with no recrudescence observed even with 10^9^ parasites (MIR >9). This suggests that despite their single-target nature, these compounds maintain a robust resistance profile under the conditions of the MIR assay. In contrast, parasites exposed to the PMX selective inhibitor WM92 at a starting *inoculum* of 10^8^ parasites showed recrudescence (MIR = 8), a result comparable to the control compound Ganaplacide (Fig 3A, Table 1).

**Figure 3.**
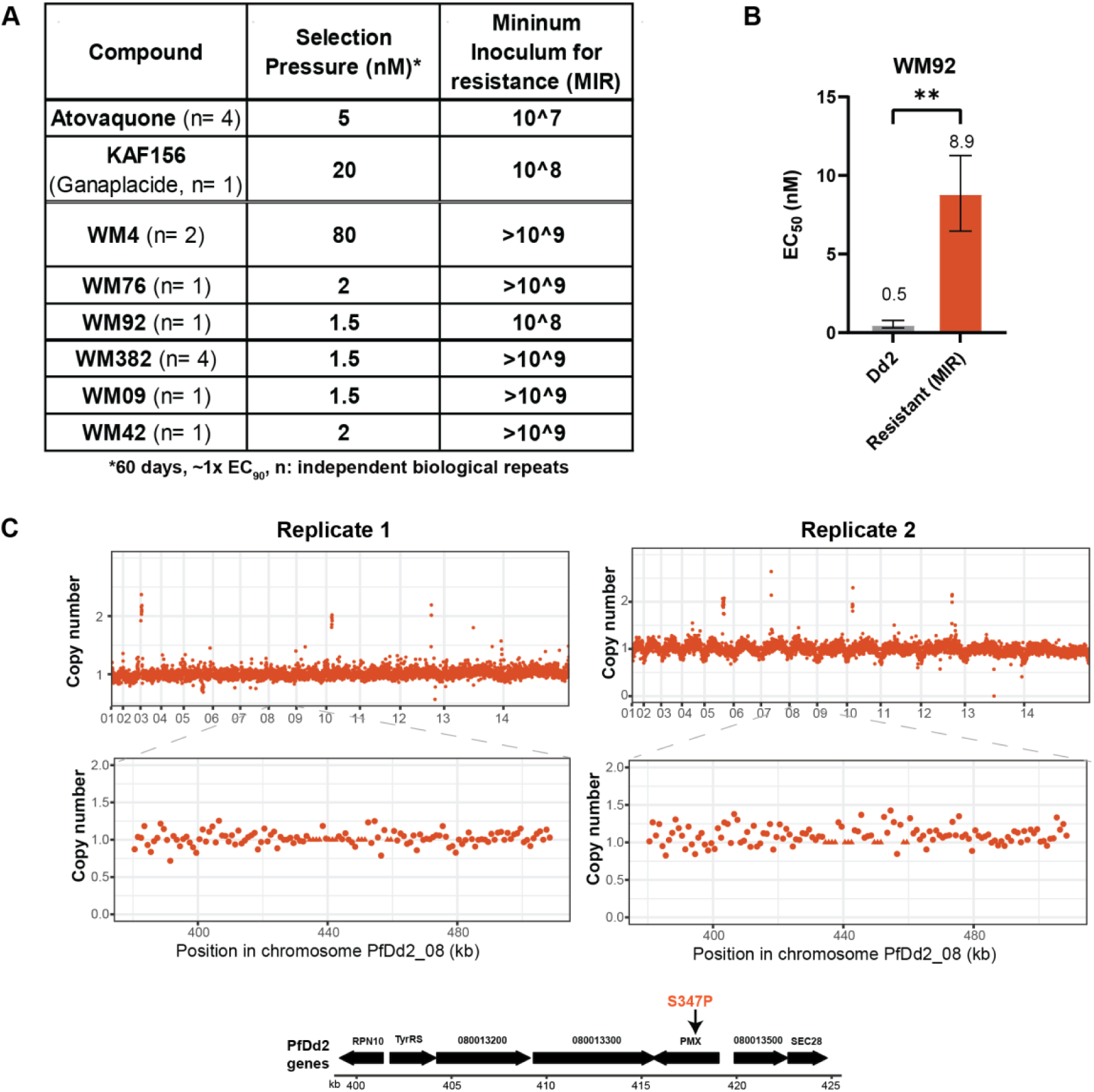
Minimum Inoculum of Resistance (MIR). **A.** Summary of risk of resistance experiments carried out with PMX-selective and PMIX/X dual inhibitors. Emergence of resistance to plasmepsin inhibitors is compared to Atovaquone and Ganaplacide. The initial *inoculum* of *P. falciparum* parasites was varied from 10^7^, 10^8^ or 10^9^ in triplicate, and resistant parasites were selected on a concentration equivalent to the ∼1x EC_90_ of selected compounds. Recrudescence of parasitaemia was followed by microscopy over 60 days for each experiment respectively. **B.** EC_50_ values (nM) derived from growth curves Dd2 wildtype and WM92-resistant parasite lines for WM92. Error bars represent the SD of >3 independent experiments. Statistical analysis conducted by Ordinary One-way ANOVA with multiple comparisons in GraphPad Prism. ** p<0.01. **C.** Genome of two replicates of *P. falciparum* WM92-resistant parasites generated from MIR studies showing whole-genome copy number (top) and detailed region of chromosome 8 (bottom). Gene numbers identified in this region of chromosome 8 can be found at PlasmoDB: https://plasmodb.org/plasmo/app/. The S347P SNP identified in the *pmx* gene is highlighted.

The decreased sensitivity of the WM92-selected parasites was confirmed with standard growth inhibition assays. The selected parasites demonstrated an 18-fold increase in EC_50_ (8.9 nM) compared to the parental Dd2 line (EC_50_ 0.5 nM) (Fig 3B). To elucidate the genetic basis of this decreased sensitivity, whole genome sequencing on two replicates from the WM92-selected resistant population was performed. Unlike the resistance mechanism observed in the incremental selection experiments (performed with *P. falciparum* with 3D7 parasites), no copy number variation of the *pmx* gene was detected in the Dd2 WM92-selected line. Instead, a single nucleotide polymorphism resulting in an S347P amino acid change in the *pmx* coding region was identified, which is equivalent to the S359P position in 3D7 parasites. This mutation likely plays a crucial role in conferring decreased sensitivity to WM92. A summary of all resistant lines generated against PMX-selective inhibitors and their corresponding genomic variants can be found in S2 and S3 Tables.

### PMX mutants decrease activity of PMX-selective inhibitors

To investigate the role of the identified SNPs and their contribution to resistance, each was introduced into the *pmx* gene of parental 3D7 parasites (S3A Fig). The presence of the expected nucleotide changes in the *pmx* gene of each genetically modified parasite lines was verified by sequencing (S3C Fig). Expression of the resulting mutant proteins were confirmed by western blot using anti-HA antibodies (S3B Fig), demonstrating that SNP insertion did not affect adversely affect PMX protein stability. All transgenic parasite strains were tested in growth inhibition assays to determine EC_50_ values for inhibition by the compounds WM92 and WM4 (Fig 4).

**Figure 4.**
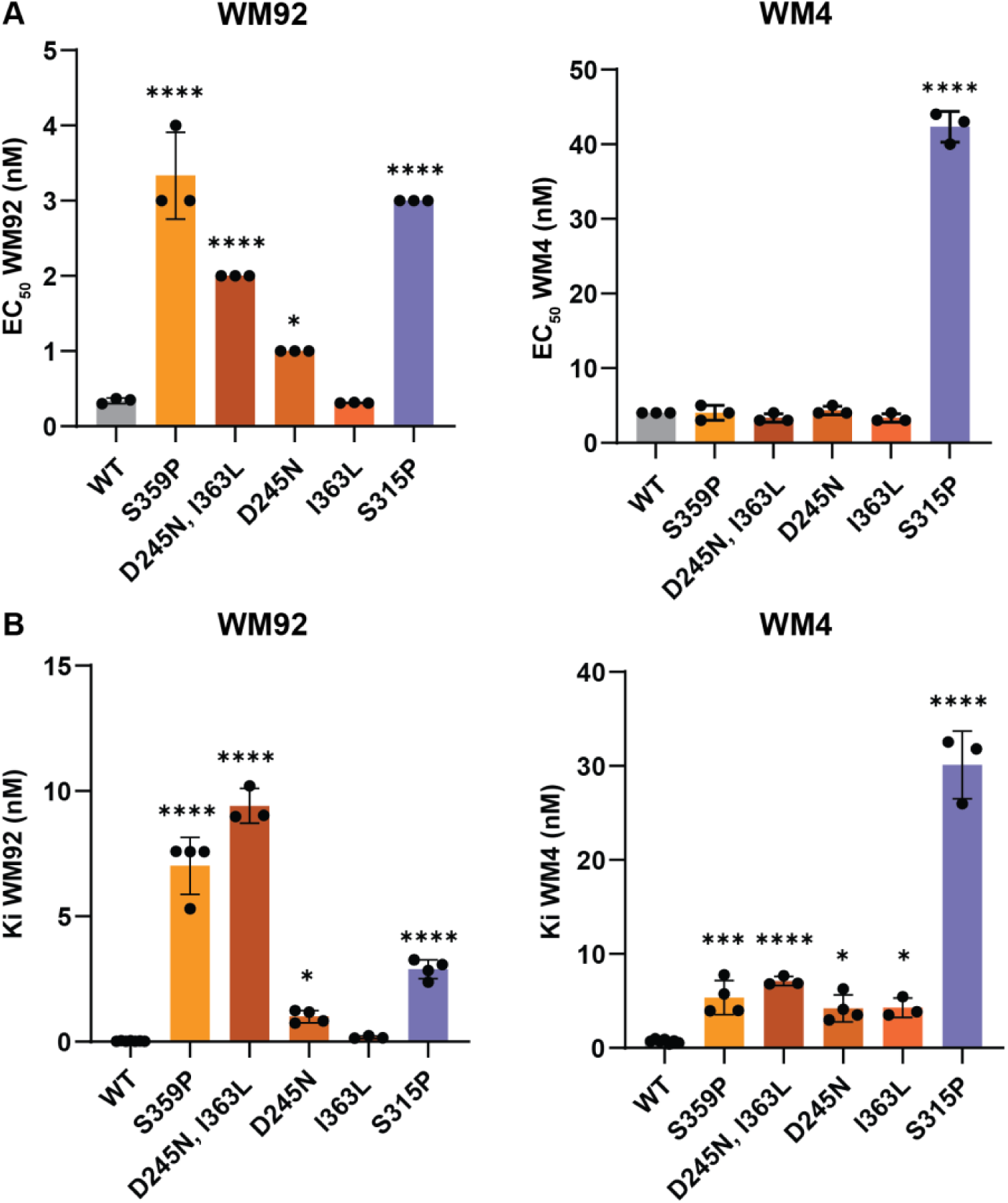
PMX mutations introduced into wildtype *P. falciparum* parasites and recombinant PMX confer resistance to WM92 and WM4. **A.** EC_50_ values for WM92 (left panel) and WM4 (right panel) were determined using growth curves of *P. falciparum* 3D7 (WT) parasites, and transfected parasite lines expressing the PMX amino acid mutations S359P, D245N/I363L, D245N, I363L or S315P. **B.** Recombinant full-length PMX protease was expressed and purified in its wildtype form (WT) and with the indicated mutations (S359P, D245N/I363L, D245N, I363L and S315P). The inhibition constant (K_i_) was determined for each protease variant. Error bars represent the SD of >3 independent experiments. Statistical analyses conducted by Ordinary One-way ANOVA with multiple comparisons in GraphPad Prism. * p<0.05, ** p<0.01, **** p<0.0001. ns = not significant.

Introduction of the S359P mutation, identified in resistant parasites selected through the MIR studies, into the PMX protease caused a significant decrease in sensitivity to the PMX selective inhibitor WM92, with the EC_50_ for parasite growth increasing approximately 10-fold from 0.3 nM to 3.3 nM (Fig 4A). The combination of D245N and I363L mutations, selected through slow incremental increase in drug concentration, in PMX expressed by the transfected parasite line also exhibited reduced sensitivity to WM92, with an EC_50_ of approximately 2 nM. Interestingly, when introduced individually into *P. falciparum*, the D245N mutation caused a modest increase in EC_50_ to approximately 1 nM, while I363L had no significant effect. This suggests an obligate co-evolution between D245N and I363L in conferring decreased sensitivity to WM92. Notably, the S359P, D245N, and I363L mutations when introduced into PMX had no substantial effect on sensitivity of the parasite to the PMX-selective inhibitor WM4, highlighting the compound-specific nature of these resistance mechanisms. In contrast, the S315P mutation, selected under WM4 drug pressure, increased the EC_50_ for both WM92 and WM4, indicating a broader resistance profile due to critical changes in the S2 loop that significantly influence the binding site for these compounds (Fig 4A).

To further analyse the effect of these amino acid changes on PMX protease activity, mutant versions of the enzyme were recombinantly expressed and characterised (Fig 4B). The S359P mutation in PMX increased the K_i_ for WM92 from 0.1 nM to 7 nM, confirming reduced affinity of binding for the compound (Fig 4B). Similarly, the combination of D245N and I363L mutations increased the K_i_ to 9 nM, while individually these mutations had minimal impact on WM92 binding affinity (Fig 4B). This combination effect at the enzyme level for K_i_ was consistent with the observed increase in EC_50_ in the parasite growth assays.

For the WM4-selected S315P mutation, an increase in K_i_ from 1 nM to 30 nM for the compound WM4 was observed, consistent with the elevated EC_50_ in parasite growth assays (Fig 4A, B). The S359P, D245N/I363L, D245N and I363L mutations caused only minor increases in K_i_ for WM4, aligning with their minimal impact on parasite sensitivity to this compound (Fig 4B). These results demonstrate that specific mutations in PMX can confer resistance to PMX-selective inhibitors through reduced binding affinity. The differential effects of these mutations on WM92 and WM4 sensitivity highlight the complexity of drug-target interactions and the potential for compound-specific resistance mechanisms.

### Modelling of compounds in PMX structure

To understand the possible ramifications of the resistant mutations on compound binding, protein models of WM92 and WM4 in complex with *P. falciparum* PMX were created (Fig 5A-E). The I363L mutation located in the S_1_’ region of PMX is likely to perturb the binding of the P_1_’ ethyl group of WM92 to this pocket (Fig 5B). The D245N and S359P mutations surround the S_3_ pocket, which houses the P_3_ chromane moiety of WM92 (Fig 5C). Although D245 and S359 do not directly interact with the P_3_ chromane moiety, mutations in these amino acids are likely to cause a conformational change in nearby amino acids that interfere with the binding of this moiety to the S_3_ pocket.

**Figure 5.**
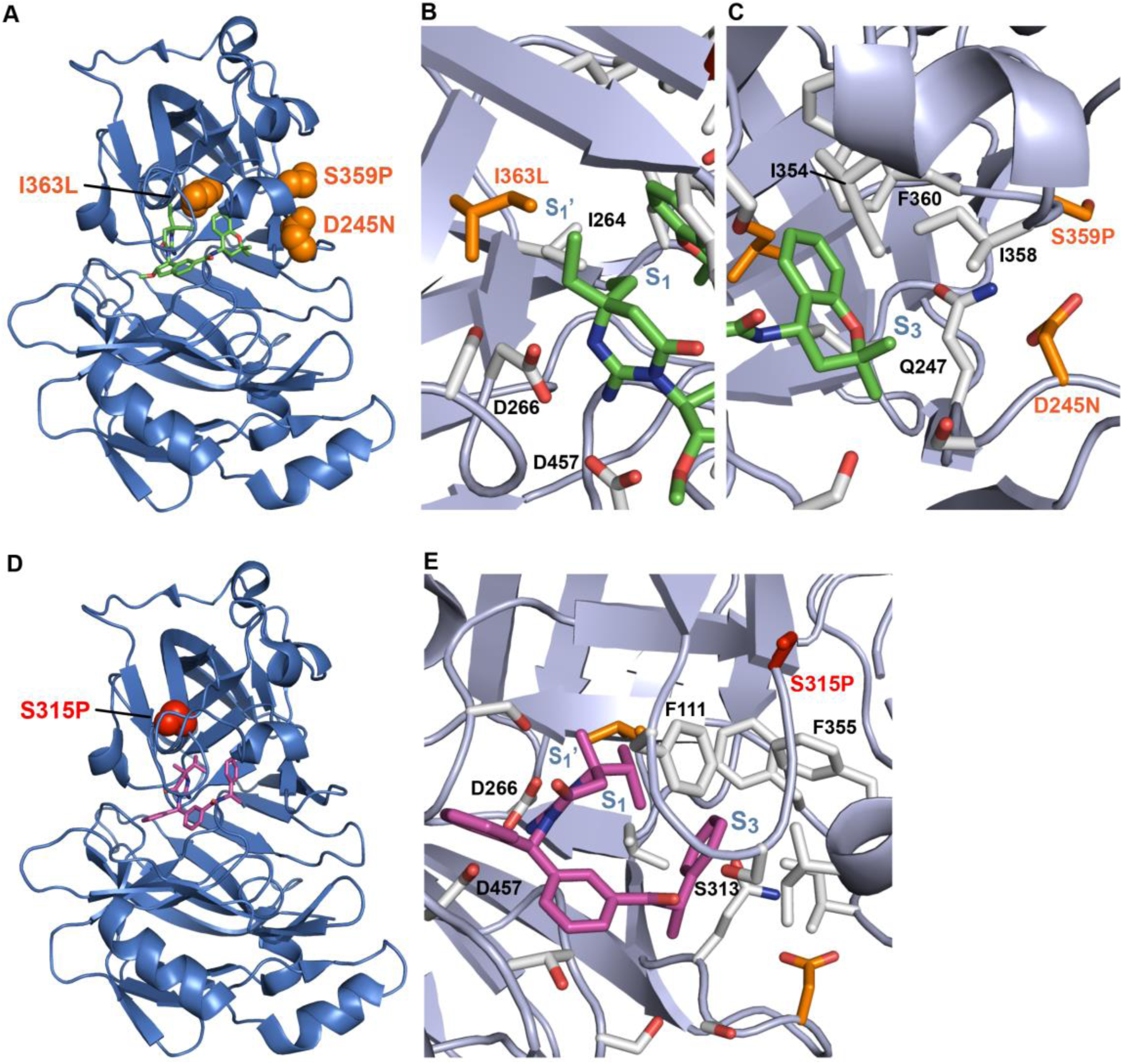
Models showing the location of the WM92 and WM4 resistant mutations in *P. falciparum* PMX. **A.** The position of the WM92 resistant mutations (orange) surrounding the binding site of WM92 (green) in PMX (blue). **B.** Close-up of the S_1_’ region surrounding the I363L mutation and the P_1_ ethyl group of WM92. **C.** Close-up of the S_3_ region showing the location of the S359P and D245N mutations in relation to the P_3_ chromane group of WM92. WM92 was modelled by building from the structure of WM382 bound to *P. falciparum* plasmepsin X (PDB 7TBC) [22]. **D.** The position of the WM4 resistant mutation S351P (red) located on the flap of plasmepsin X on top of the binding site of WM4. **E.** Close-up of the flap region showing the binding orientation of WM4. WM4 was modelled by superimposing the structure of *P. vivax* PMX in complex with WM4 (PDB 7TBE) with the structure of WM382 bound to *P. falciparum* PMX (PDB 7TBC) [22].

Critically, S315 is located on the flap and does not directly interact with WM4 (Fig 5D, E), yet a change from Ser to Pro causes a conformational change in the flap structure that perturbs amino acids directly interacting with WM4, reducing its ability to bind to PMX. Notably, S315 represents a non-essential residue in the substrate binding architecture; however, its mutation to P315 causes an increase in Ki of PMX for WM4 indicating that this change decreases the affinity of binding of this compound resulting in an increase in EC_50_ for *P. falciparum* growth assays (Fig 4A, B). However, it appears the parasite can tolerate alterations to S315 because it does not participate in the essential interactions required for substrate recognition and proteolytic activity. Conversely, the parasite cannot afford to mutate residues critical for PMX function without compromising parasite replication fitness. Thus, the mutations conferring resistance to WM4 exploit regions of the protease that are functionally malleable for substrate binding, allowing decrease inhibitor binding, while preserving PMX catalytic competence.

### Fitness cost of pmx gene amplification and mutations in P. falciparum

To assess the potential fitness costs associated with *pmx* gene amplification and mutations in the aspartic protease, growth competition assays were performed. These assays involved mixing equal number of wildtype and resistant parasites, culturing them over 28 days, and measuring EC_50_ values at the start and at the end of the experiment (Fig 6A). When cultured individually, the WM92-resistant (10x *pmx* amplification and D245N/I363L mutations), WM92-revertant (2x *pmx* amplification and D245N/I363L mutations), and parental 3D7 strains maintained stable EC_50_ values throughout the 28-day period, as expected. However, in direct competition, both WM92-resistant and WM92-revertant parasite lines showed significant fitness disadvantages compared to the parental 3D7. When WM92-resistant parasites were mixed with an equal number of 3D7 parasites, the initial EC_50_ of >10 nM decreased substantially to 0.4 nM by day 28, approaching that of 3D7 alone (Fig 6A). Similar results were observed for the WM92-revertant strain. Notably, the similar outcomes for WM92-resistant and WM92-revertant in the competition assays suggest that the fitness cost of the mutations in PMX(I363L/D245N) was significant compared to that of the gene amplification, as the revertant strain maintains the mutations but has reduced copy number.

**Figure 6.**
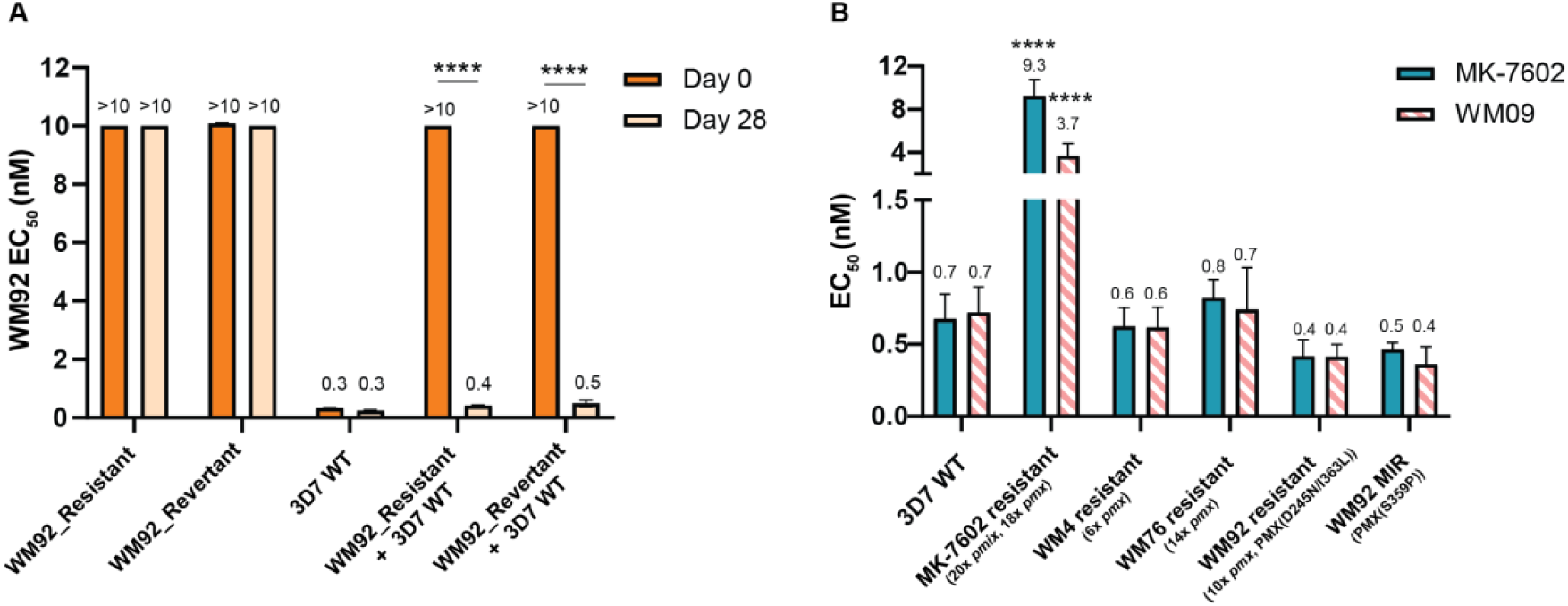
PMX mutant parasites are associated with a reduced parasite fitness and remain sensitive to dual PMIX/X inhibitors. **A.** WM92 resistant and reversion lines containing amplifications and mutations in *pmx* were mixed 1:1 with 3D7 wildtype (WT) parasites and the EC_50_ evaluated using LDH growth assays on Day 0 and 28, alongside individual counterpart lines. Error bars indicate the SD of two technical replicates. Statistical analysis was performed in GraphPad Prism via a two-way ANOVA with multiple comparisons. **** indicates p<0.0001. ns= not significant. **B.** EC_50_ values depicted for dual PMIX/X inhibitors, WM09 and clinical candidate MK-7602, against the PMX mutant lines showing sensitivity of the dual inhibitors is maintained. Error bars indicate the SD of >3 biological replicates. Statistical analysis was performed in GraphPad Prism via a one-way ANOVA with comparison to 3D7 WT. **** indicates p<0.0001. No asterisks indicate no significance.

The same fitness cost experiment was repeated with the mutant PMX parasite lines including PMX(D245N), PMX(I363L) and PMX(D245N, I363L) using PMX(WT) as the benchmark for normal parasite growth (S4 Fig). This direct growth competition experiment showed that PMX(D245N) and PMX(D245N, I363L) had moderate levels of fitness cost with a 2.3-fold and 2.7-fold reduction in EC_50_, respectively, after 28 days from being mixed 1:1 with PMX(WT) parasites (S4 Fig). In contrast, PMX(I363L) alone showed no fitness cost with equal EC_50_ values observed on day 0 and 28 (EC_50_ = 0.2 nM) (S4 Fig). However, as PMX(I363L) exhibits an EC_50_ comparable to wildtype it may still be outcompeted, though any difference would likely be below the level of detection. These findings demonstrated that the genetic changes in *pmx*, including both gene amplification and mutations, impose a fitness cost on the parasite, with the greatest impact observed with gene amplification of *pmx.*

The relationship between the resistance mechanisms selected by PMX-selective compounds (WM4, WM76, and WM92) and dual PMIX and X inhibitors WM09 and MK-7602 were further investigated. Accordingly, growth inhibition assays were conducted to evaluate potential cross-resistance between parasite lines containing variants in PMX and dual PMIX and X inhibitors. Importantly this revealed that the WM4-resistant (6x *pmx*), WM76-resistant (14x *pmx*), and WM92-resistant (10x *pmx*, D245N, I363L) selected lines all remained sensitive to both MK-7602 and WM09 dual inhibitors (Fig 6B, S4 Table). In contrast, a previously described MK-7602 selected parasite line with amplified copies of both *pmx* and *pmix* [19] showed shifts in EC_50_ values compared to a 3D7 line of 15-fold and 4-fold against MK-7602 and WM09, respectively.

A particularly noteworthy observation was the differential response of the WM4-resistant line to PMX-selective versus PMIX/X dual inhibitors. This strain, characterized by increased PfPMX expression, demonstrated a 6-fold increase in EC_50_ for WM4 (66.6 nM compared to 11 nM in 3D7 WT), a 2.7-fold increase for WM92 (2.25 nM compared to 0.82 nM in 3D7 WT) and 4.1-fold increase for WM76 (3.5 nM compared to 0.845 nM in 3D7 WT) (Table S4). However, the PMIX/X dual inhibitors WM382, WM09 and MK-7602 maintained their potency against this line, with EC_50_ values (∼0.6 nM) nearly identical to those observed in the parental 3D7 strain (0.5 nM, 0.7 nM and 0.7 nM, respectively). The fact that dual inhibitors retained their efficacy against the WM4-resistant line underscores the advantage of targeting both PMIX and PMX simultaneously, as it mitigates the impact of PMX overexpression as a resistance mechanism.

This comparison highlights the distinct resistance profiles of PMX-selective and dual PMIX/X inhibitors and underscore two critical points. Firstly, the genetic changes conferring resistance to PMX-selective inhibitors come at a significant fitness cost to the parasite, potentially limiting the spread of such resistant strains in the absence of drug pressure. Secondly, dual PMIX and X inhibitors maintain their efficacy against parasites with PMX-specific resistance mechanisms, demonstrating both the importance of PMIX in their mode of action and their high barrier to resistance. This robustness persists even in the face of substantial genetic alterations to PMX, reinforcing the potential of dual-target inhibition as a strategy for antimalarial drug development.

## Discussion

The emergence and spread of antimalarial drug resistance poses a major threat to global malaria control efforts. In this context, our data provide important insights into the resistance profiles of PMX-selective versus dual PMIX/X inhibitors and highlight the potential advantages of multi-target inhibition [17, 19]. Both inhibitor classes exhibit a relatively high barrier to resistance; however, the dual PMIX/X inhibitors display a clearly superior profile. Even after prolonged *in vitro* drug pressure, we were unable to select parasites with stable resistance to the dual inhibitors, whereas resistance to PMX-selective compounds could be generated, although only with considerable difficulty. Previous work on PMX-selective series has shown that resistance to the compound UCB7362 can be obtained at a minimum *inoculum* (MIR) of 9.6 × 10^6^, and that a related analogue (compound 3) can select resistance at an *inoculum* of 5.3 × 10^5^, albeit with only a modest EC_50_ shift between parental and mutant lines [32]. These findings suggest that the ease of resistance selection is influenced not only by the essentiality of PMX, but also by features of the compound scaffold.

When placed in the broader context of antimalarial development, the resistance profile of dual PMIX/X inhibitors appears particularly robust. For several clinical-stage candidates, including Ganaplacide, Cipargamin, and DSM265, resistant parasites have been reported to emerge within 3–4 months of continuous drug pressure [12, 24, 33]. The acylguanidine MMV688533 shows a lower propensity for resistance, yet mutations in PfACG1 and PfEHD still confer measurable reductions in potency [34]. By contrast, under comparable or more prolonged *in vitro* selection conditions, we were unable to derive stable resistant lines to dual PMIX/X inhibitors, indicating a substantially higher barrier to resistance. This exceptional profile supports further development of dual PMIX/X inhibitors as promising, durable components of future combination regimens aimed at delaying or preventing the emergence of clinical resistance.

This robustness is particularly noteworthy when compared to the resistance profiles of current front-line antimalarials. Artemisinin partial resistance, mediated primarily through mutations in the K13 propeller domain, has emerged and spread relatively quickly in Southeast Asia [4]. Similarly, resistance to partner drugs such as piperaquine has developed through amplification of *plasmepsin 2* and *3* genes and mutations in PfCRT [7, 35]. Importantly, the Chloroquine-resistant PH1263-C parasite line, which carries multiple mutations including a *pmII* amplification, remained fully susceptible to all tested PMX-and PMIX/X dual-inhibitors (S4 Table). The high barrier to resistance observed with dual PMIX/X inhibitors is more comparable to that seen with some HIV antiretrovirals, such as second-generation integrase strand transfer inhibitors like dolutegravir [36].

A critical conclusion from this study is that PMX-selective compounds should not be prioritised as antimalarial agents. Although they are potent and well-behaved chemically, their clinical use would focus selection pressure solely on PMX and risk undermining the major advantage of dual PMIX/X inhibitors and an exceptionally high barrier to resistance. We show that PMX-selective compounds can drive both *pmx* gene amplification and the emergence of point mutations in the PMX protease. These adaptations enable parasites to tolerate otherwise inhibitory PMX-targeted pressure. Importantly, however, the resistance phenotypes are not uniform across the PMX-selective series: individual mutations confer resistance to the selecting compound but only partial, or no resistance to other PMX-selective inhibitors. This pattern was consistent with subtle, chemotype-specific differences in how individual inhibitors engage the PMX active site and how particular amino acid substitutions perturb those interactions. Nevertheless, from a population perspective, widespread use of any PMX-selective agent would be expected to seed a diversity of *pmx* alleles, through amplification and mutation, that could collectively lower the evolutionary barrier to resistance across the class and partially diminish the robustness of dual PMIX/X inhibitors by weakening one arm of their dual-target mechanism. Notably, even in the presence of PMX variants and *pmx* amplification, dual PMIX/X inhibitors retain high activity, because they also target the essential protease PMIX and multiple essential PMIX-dependent processes. This underscores the intrinsic resilience of dual-target inhibition and supports a deployment strategy that prioritises dual PMIX/X inhibitors while avoiding clinical use of PMX-selective agents.

The mechanisms of resistance to PMX-selective inhibitors involved both gene amplification and point mutations in the PMX protease. This finding aligns with previous observations of gene amplification as a resistance mechanism in *Plasmodium*, such as the amplification of *pfmdr1* in mefloquine resistance [29, 37] and plasmepsin 2 and 3 in piperaquine resistance [7]. The identification of specific point mutations (S315P, D245N, I363L, S359P) provides valuable insights into the molecular basis of drug-target interactions and resistance development. Interestingly, our results revealed compound-specific resistance profiles even among structurally related PMX-selective inhibitors. This highlights the complexity of drug-target interactions and emphasizes the need for careful consideration of resistance profiles during drug development.

The structural modelling of PMX in complex with the inhibitors provides insights into the molecular mechanisms of resistance and offers guidance for future drug design strategies [22]. The identified mutations appear to affect inhibitor binding through various mechanisms, including steric hindrance, conformational changes, and alterations in pocket geometry. These findings align with previous studies on protease inhibitor resistance in other systems, such as HIV protease inhibitors [38].

The fitness cost associated with PMX amplification and mutations is a crucial finding of the study. Competition assays demonstrated that resistant parasites were significantly outcompeted by wild-type parasites in the absence of drug pressure. This fitness cost could potentially limit the spread of resistant parasites in natural populations [39]. The cross-resistance studies, including the Antimalarial Resistome Barcoding (AReBar) assays, revealed no pre-existing resistance to either PMX-selective or PMIX/X dual inhibitors among known resistant *P. falciparum* strains. This lack of cross-resistance with existing antimalarials shows that these compounds could be effective against current drug-resistant parasite populations [9].

The inability to design PMIX-selective inhibitors during the medicinal chemistry program was noteworthy and likely reflects fundamental differences in the structural architecture of the S’ binding sites between PMIX and PMX. The large loop structure in PfPMX, and its associated conformational mobility, likely creates a more permissive S’ pocket that tolerates substantial structural variation. Conversely, dual PMIX/X inhibitors achieve their broad efficacy by concentrating their interactions within the S pockets - regions that are typically more critical for substrate binding and thus more structurally conserved across both proteases.

In conclusion, our findings strongly support the development of dual PMIX/X inhibitors as promising antimalarial candidates with a high barrier to resistance. The superior resistance profile of these dual inhibitors, combined with their efficacy against existing drug-resistant strains, positions them as valuable tools in the ongoing fight against malaria. As we move forward, it will be essential to focus on dual inhibitor development, continue monitoring for potential resistance mechanisms, and explore combination therapies that could further enhance the longevity and efficacy of these promising antimalarial compounds [9].

## Materials and Methods

### Ethics Statement

Use of human blood and serum was approved by the Walter and Eliza Hall Institute of Medical Research Human Ethics committee under approval number 23-05VIC-09.

### Plasmodium falciparum Culture

*P. falciparum* asexual blood stage parasite cultures and the parasite lines derived from these by genetic manipulation were grown in *in vitro* culture [40]. All parasites were supplied with O^+^ erythrocyte (Australian Red Cross Bloodbank, South Melbourne, Australia) at 4% hematocrit in Roswell Park Memorial Institute (RPMI) 1640 medium supplemented with 26 mM 4-(2-hydroxyethyl)piperazine-1-ethanesulfonic acid (HEPES), 50 μg/ml hypoxanthine, 20 μg/ml gentamicin, 2.9% NaHCO_3_, and 5% Albumax II^TM^(Gibco) + 5% heat-inactivated human serum or 10% Albumax II^TM^ (Gibco). Cultures were incubated at 37 °C in a gaseous mix of 94% N_2_, 1% O_2_ and 5% CO_2_. Parasites were sub-cultured with new culture media every 2 days to maintain 4% haematocrit and keep the parasitemia below 5%.

### Construction of P. falciparum lines expressing HA-tagged proteins

Transgenic parasite lines were made using the CRISPR-cas9 system as previously described [17]. This involved generation of a guide plasmid and a homology directed repair (HDR) plasmid that replaces the endogenous target gene with a version carrying the desired SNPs and three haemagglutinin tags for validation of gene targeting. The guide plasmid was created by InFusion cloning of the guide oligos MD1/MD2 into *Btg*ZI restriction enzyme digested pUF1-Cas9G vector. These oligos induce a Cas9 dependent double-stranded break at position 496 of the PfPMX protein coding sequence. A modified p1.2 based plasmid encoding WR99210 resistance was used for HDR plasmid creation. Firstly, a 532 bp 3’ homology arm of PfPMX was PCR amplified from *P. falciparum* 3D7 strain genomic DNA using the oligos MD3/MD4 and inserted into the vector using the restriction enzymes *EcoRI* and *Pst*I. This intermediate vector was subsequently modified by inserting the synthetic gene fragment consisting of a combined 5′ homology arm and codon-optimised sequence, containing the desired gene mutations and epitope tag (Genscript), using the restriction enzymes *Not*I and *Xho*I. 50 mg of linearized HDR plasmid and 100 mg of circular guide plasmid were transfected simultaneously into *P. falciparum* parasites that had been previously synchronised using 5% sorbitol and purified using a Percoll gradient to purify schizont stages. This was performed by layering cultured, synchronized *P. falciparum*-infected red blood cells over a pre-formed 35%/65% Percoll step gradient, followed by centrifugation (1,500 × *g*, 15 min, no brake) to separate mature, lower-density schizonts at the interface. Parasites with an integrated drug resistance cassette were selected and maintained on 2.5 nM WR99210. Integration was confirmed via PCR, Sanger sequencing (AGRF) and anti-HA-HRP (1:2000, ThermoFisher) western blotting.

### P. falciparum growth inhibition assays and EC_50_ determination

The *P. falciparum* growth inhibition assay was conducted with minor modifications [41]. An inoculum of synchronized ring-stage parasitized red blood cells (pRBC) at 0.7% parasitemia and 2% haematocrit in complete media was used for the assay. Assay plates (Greiner #781098, 384 well, white, tissue culture treated) in 16-point dilution series of the compounds were prepared in assay plates using an Echo555 (Labcyte). Appropriate volumes of 10 mM compound stocks were transferred into the assay plates in duplicates, such that the starting concentration was 11.25 µM, with a 1:3-fold dilution series. All wells were backfilled with DMSO to 45 nL such that this remained constant (0.1% DMSO). The growth control was 0.1% DMSO and the negative growth control was 2.5 µM chloroquine. A parasite inoculum (40 μL) was dispensed into plates containing compounds. Following incubation for 72 hours, plates were sealed with parafilm and frozen at -80°C overnight. Plates were thawed at room temperature for at least 4 hours prior to lactase dehydrogenase (LDH) activity being measured. 45 μL of fresh LDH reaction mix (174 mM sodium l-lactate, 214 µM 3-acetyl pyridine adenine dinucleotide (APAD), 270 µM Nitro Blue tetrazolium chloride (NBT), 4.35 U ml^-1^ diaphorase, 0.7% Tween 20, 100 mM Tris-HCl pH 7.5) was dispensed into each well using a Multidrop Combi dispenser at high speed to allow good mixing. Absorbance at 650 nm was measured in an EnVision (PerkinElmer) plate reader after 20 min of incubation at room temperature. Data were normalized to percent viability using positive and negative controls. EC_50_ values (relative inflection of dose response curve) were calculated by Dotmatics 5.3 and Spotfire 7.11.1 software using a nonlinear regression four-parameter fit analysis. The equation used is sigmoidal dose response (variable slope), Y = Bottom + (Top – Bottom) / (1 + (EC_50_ / X)^HillSlope^).

For all other experiments, the determination of EC_50_ values was obtained using ring stage parasites at 0.5% parasitemia which were dispensed in a 50 µL culture at 2% haematocrit in 96 well round bottom microtiter plates (Falcon) with doubling dilutions of each compound. After 72 hours incubation at 37°C each well was fixed at room temperature for 30 min with 50 µL of 0.25% glutaraldehyde (Pro-SciTech) diluted in PBS. Following centrifugation at 1200 rpm for 1 min, supernatants were discarded, and parasites were stained with 50 µL of 2.5-5X SYBR Green (Invitrogen) diluted in PBS. 50,000 cells were counted by flow cytometry using a Attune Nxt Flow Cytometer (ThermoFisher) or Cell Lab Quanta SC - MPL Flow Cytometer (Beckman Coulter). Growth was expressed as a percentage of the parasitemia obtained using a vehicle control. All samples were tested in triplicate. EC_50_ values were determined using GraphPad Prism.

### Expression and purification of PfPMX, PfPMIX and PfPMX mutants

The apo form of PfPMX wild type (T27 to N573) and the corresponding single mutants PfPMX S315P, PfPMX S359P, PfPMX D245N, PfPMX I363L and the double mutant PfPMX D245N/I363L were expressed in HEK expi 293 cells, using the pTriex2 vector with the inserted gene sequences codon optimised for expression in mammalian cells (Bioneer) [17]. Sequence corresponding to the TEV-FLAG peptide was inserted 5’ to the stop codon to assist with the purification of the expression product and removal of the FLAG peptide tag. Expression of the various PfPMX proteins was conducted as per the manufacturer’s protocols. Culture supernatants were harvested at day 4 of culture (viable cell density of 72%). PfPMX proteins were extracted from the supernatant using M2 antiflag affinity resin (Sigma). Bound protein was eluted using 3 column volumes of 0.1 mg/ml flag peptide. Fractions containing the desired expression product were pooled then concentrated and further purified using a Superdex 200 10/30 size exclusion column (Cytiva) run in the presence of 20mM Tris pH7.2/150mM NaCl. The eluted fractions were analysed using SDS-PAGE and those containing the highly purified PfPMX proteins were pooled for further studies. The produced versions of PfPMX had similar autolytic products to those observed by previously [17, 42].

PfPMIX was expressed as previously described [22]. Briefly, to express recombinant PfPMIX the gene sequence (37KDC.NNL627) was recodoned for insect cell expression and synthesised by GenScript and subcloned into the p1TF vector that enabled expression in insect cells. PfPMIX was expressed in CHO cells as described [43]. PfPMIX was purified by nickel affinity chromatography and ion exchange chromatography. The PfPMIX was flash frozen and stored at 80°C until further use.

### Substrate peptide Km determination

The K_m_ of Rh2N (DABCYL-HSFIQEGKEE-EDANS synthesized by Chinapeptides) was determined by titrating the fluorogenic peptide in 12-point 2-fold dilution series (0.08-80 µM) against each PMX enzyme at their optimal concentration (0.25 to 2.5 nM depending on enzyme activity). All reagents were prepared in buffer containing 25 mM sodium acetate (pH 5.5) and 0.005% Tween-20. The assay was conducted in 20 µl volume in a 384 well black plate (Corning #3820) with each PMX enzyme reaction initiated by the addition of 10 µl of PMX enzyme to plate containing 10 µl of titrated Rh2N. The enzyme kinetic assay was allowed to proceed at 37 °C for 30 minutes in the PHERAstarFSX plate reader (BMG Labtech). Samples were excited at 340 nm and fluorescence emission measured at 480 nm at one minute interval. Linear slope (0-15min) was used to determine the initial velocity (RFU/min) at each Rh2N concentration and plotted in Graphpad Prism to calculate the substrate Km based on Michaelis-Menten enzyme kinetics. Reported Km values were calculated on the Mean ± SD of three independent dataset.

### Plasmepsin X fluorogenic assay

Compound potency assay against each PMX enzyme was conducted in 15 µl reaction volume. Prior to that each PMX enzyme was titrated at 10 µM Rh2N in a time-course manner to determine the reaction linearity and the optimal protein concentration (in bracket below) for use in compound testing. Compounds (synthesized by Wuxi) were plated with 16-point 3-fold dilution series starting at 10 µM in triplicates in the 384 well black plates (Corning #3820) using an Echo acoustic dispenser (Labcyte) with all wells backfilled to 150 nl DMSO (1% final) and repeated in three independent experiments. Compounds were pre-incubated with 7.5 µl PMX enzyme (0.1 nM PMX, 0.1 nM PMXS315P, 0.1 nM PMXD245N, 0.2 nM PMXS359P, 0.05 nM PMXI363L, 0.025 nM PMXD245N/ PMXI363L) at room temperature for 20 mins and the reaction initiated with 7.5 µl Rh2N (10 µM final). Plate was incubated in a humidified incubator at 37 °C for 4 hours. Fluorescence output was measured in a PHERAstar Microplate Reader (BMG LABTECH) at excitation 340 nm and emission 480 nm. Data were normalized to percent inhibition relative to 1% DMSO (high control) and 1 µM WM382 (low controls) and plotted in Graphpad Prism. IC_50_ values were calculated using a nonlinear regression four-parameter fit analysis. The equation used is sigmoidal dose response (variable slope), Y = Bottom + (Top – Bottom) / (1 + (IC_50_ / X)^HillSlope^). Compound Ki values were calculated from the IC_50_ values using the Cheng-Prusoff equation, Ki = IC_50_/(1 + [S]/Km). Reported Ki values were calculated on the Mean ± SD of three independent dataset.

### PMX and PMIX primary screening method

PMX and PMIX primary screening were conducted in 1536 well plate (black low volume non-binding plate, Corning #3728) at a single compound dose of 10 nM in duplicates in 5 µl assay volume using in-house produced PfPMX (0.2nM) or PfPMIX (5 nM) recombinant proteins. 1536 well assay ready plates were prepared using an Echo acoustic dispenser (Labcyte) with 5nl of 10 µM compound (1% DMSO) transferred into the 1536 well plate. 2.5 µl recombinant protein (PMX or PMIX) was dispensed into compound containing assay plates. Plates were allowed to incubate for 20 min at room temperature, and the reaction was started with a further 2.5 µl addition of 1.8 µM PMX substrate (i.e. Rh2N, DABCYL-HSFIQEGKEE-EDANS) or 7.2 µM PMIX substrate (i.e. RON3, DABCYL-KEISFLERRE-EDANS). Reaction was incubated at 37 °C for 2 hours. Enzyme and substrate working solutions were prepared in assay buffer consisting of 25 mM sodium acetate, 0.005% Tween-20, pH 5.5 for PMX assay and same buffer plus 100 mM sodium chloride for PMIX assay. Samples were excited at 340 nm and fluorescence emission measured at 480 nm using a PHERAstarFSX plate reader (BMG Labtech). Data were normalized to percent inhibition using positive and negative controls. The 0% inhibition control contained 1% DMSO and the 100% inhibition control contained 1 µM WM382. 931 hits were selected for activity confirmation in 5-points/10-fold dilution series in the same assays. IC_50_ values (relative inflection of dose response curve) were calculated by Dotmatics 5.3 and Spotfire 7.11.1 software using a nonlinear regression four-parameter fit analysis. The equation used is sigmoidal dose response (variable slope), Y = Bottom + (Top – Bottom) / (1 + (IC_50_ / X)^HillSlope^).

### In vitro selection of resistant parasites and reversion line

Three replicate cultures of clonal *P. falciparum* 3D7 parasites were grown on incremental increases of compound beginning at a concentration of 2x EC_50_. When parasitemia was significantly reduced, compound pressure was removed and when parasitemia recovered, compound pressure was resumed. This was repeated until the parasites were adapted to higher drug concentration over the course of up to 15 months. For the reversion line, drug pressure was removed from the resistant lines and cultures were maintained in drug-free complete media for 6 months. EC_50_ values were determined from these lines and genomic DNA was purified (QIAGEN Blood and Tissue Kit) for WGS analysis.

### Genome sequencing and bioinformatics analysis

An input of 100 ng of *P. falciparum* genomic DNA was prepared and indexed for Illumina sequencing using the TruSeq DNA sample Prep Kit (Illumina) as per manufacturer’s instruction. The library was quantified using the 4200 TapeStation System (Agilent). The indexed libraries were pooled and diluted to 750 pM for paired end sequencing (2x 150 bp reads) on a NextSeq 2000 instrument using the P2 300 cycle kit (Illumina) as per manufacturer’s instructions.

The quality of sequencing was confirmed using FastQC. Fastq files were aligned to reference PlasmoDB-52_Pfalciparum3D7 using bwa-mem version v0.7.17. Sequences were filtered using Picard MarkDuplicates. Copy number analysis was performed using the R package QDNAseq v1.28.0 [44]. Structural variant calling was performed using GRIDSS v2.12.1 [45, 46]. Regions of interest were inspected with Integrated Genome Viewer. Copy number bins in QDNASeq were filtered to exclude telomere and centromere regions, and by regions with mapability less than 50% using 30mers generated by GenMap [47]. Calling of single nucleotide variants (SNVs) and indels was performed with bcftools v1.13 and calls were filtered using criteria: QUAL > 50; not an indel with length > 5 of A and T only; not in telomere, centromere or hypervariable regions [48].

Whole genome sequence data for this study have been deposited in the European Nucleotide Archive (ENA) at EMBL-EBI under accession numbers PRJEB36069, PRJEB58099, PRJEB53576 and PRJEB107760.

### Risk of resistance and minimum inoculum for resistance (MIR)

Cultures of 10^7^, 10^8^ and 10^9^ *P. falciparum* Dd2 parasites were exposed to ∼1x EC_90_ (WM382: 1.5 nM, WM09: 1.5 nM, WM4: 80 nM, WM76: 2 nM, WM92: 1.5 nM, Atovaquone: 5nM, KAF156: 20 nM) of constant drug pressure and monitored for parasite recrudescence for 60 days. Experiments were carried out in triplicate and monitored by weekly microscopic examination of thin blood films. Media and compound were replaced three times each week. Evaluation of EC_50_ and genome analysis from WM92-treated recrudescing parasites was carried out as outlined above.

### Antimalarial resistome barcoding assay (AReBar)

A parasite pool consisting of 52 barcoded lines (see Supp. Table 1) covering a range of mutant drug targets and resistance mechanisms was used to profile compounds for cross-resistance, essentially as described [19, 26]. Briefly, triplicate cultures of the pool were exposed to 3×EC_50_ of each compound for 14 days, with parasite growth measure by flow cytometry (stained with 1×SYBR Green and 200nM Mitotracker Deep Red). A drug-free control and treatment with the ATP4 inhibitor KAE609 were also included. The relative abundance of each line was measured by amplicon sequencing (Illumina platform) of the barcode region. Due to the absence of growth for cultures treated with WM382, WM4, WM92, WM76, WM09 or WM42, insufficient sample was obtained to generate an amplicon, in contrast to the drug-free and KAE609-treated parasite cultures.

### P. falciparum fitness cost assays

Synchronized ring-stage WM92_Resistant and WM92_Reversion parasite lines were mixed 1:1 with a 3D7 WT line. Synchronized ring-stage PMX(D245N), PMX(I363L) and PMX(D245N, I363L) lines were mixed 1:1 with a PMX(WT)-HA parental line. The fitness cost of the copy number increase of *pmx* and PMX SNVs were then evaluated as previously described by lactate dehydrogenase growth assays [19] with a starting titration series from 10 nM of WM92.

## Acknowledgements

The authors thank Australian Red Cross Blood Service for blood, David A. Fidock for providing the PH1263-C *P. falciparum* parasites and Sergio Wittlin for providing the Dd2-NITD609 and Dd2-SJ557733 parasites. This work was supported by The Wellcome Trust (109662/Z/15/Z and 202749/Z/16/Z), Victorian State Government Operational Infrastructure Support, Australian Government NHMRC IRIISS, and National Health and Medical Research Council of Australia.

## Lead contact and materials availability

Further information and requests for resources and reagents should be directed to and will be fulfilled by the Lead Contact, Alan F. Cowman. All unique/stable reagents generated in this study are available from the Lead Contact with a completed Materials Transfer Agreement.

## Author contributions

PF and MD designed, performed and interpreted all drug selection experiments, and analysed parasite data. JT selected parasites with WM4. WS and DM designed constructs and obtained parasite transfectants. ANH expressed and purified recombinant proteins. AN analysed kinetics of recombinant proteases, performed the PMIX and X screen and *P. falciparum* growth inhibition assays. JSP and TP analysed genomes. RC and MCSL devised and performed the AReBar assay. KL provided oversight and management of the PMIX and PMX screen. BES did the PMX structural modelling. DBO designed experiments and with MdLR and JAM designed compounds. AFC designed and interpreted experiments and with PF and MD, wrote the final manuscript. All authors read and edited the manuscript.

## Competing interests

The authors have no conflict of interest to declare.

**S1 Fig. Confirmation of PMX(D245N) and PMX(I363L) in WM92-resistant and reversion parasites by PCR.** The *pmx* gene was amplified by PCR (left) and sequenced (right), showing the presence of both mutations in WM92-resistant and reversion parasites.

**S2 Fig. Long-term selection of parasites with WM4 results in a copy number increase and a SNV in PMX. A.** Wildtype parasites were exposed to increasing amounts of WM4 up to 250 nM over two years, resulting in a EC_50_ shift of 156-fold. **B-C.** Whole genome sequencing revealed a 14x copy number amplification in a region encompassing *pmx*. **C.** A single nucleotide variant was also found in PMX, resulting in PMX(S315P). A representative clone is shown in panel C. Error bars in panel A represent the standard deviation of two biological replicates, each with three technical replicates. Statistical analysis performed via an unpaired t-test with * indicating p<0.05.

**S3 Fig. Generation of mutant parasite lines. A.** CRISPR Cas9 cloning scheme to introduce the D245N, S315P, S359P, I363L and D245N/I363L mutations into *P. falciparum* PMX. **B.** Confirmation western blot demonstrating the presence of Hemagglutinin (HA) tags on the amended parasite lines. **C.** 3’ Integration PCRs of the amended *pmx* gene. Sequenced PCR products showed single base-pair changes were present resulting in desired mutations in *pmx* coding sequence.

**S4 Fig. Mutations PMX(D245N) and PMX(D245N/I363L) but not PMX(I363L) alone results in parasite fitness cost.** Parasite lines with introduced PMX mutations were mixed 50:50 with 3D7 wildtype (WT) parasites and EC_50_’s were evaluated by lactate dehydrogenase assays (LDH) on Day 0 and 28, alongside individual counterpart lines. Error bars indicate the SD of two technical replicates. Statistical analysis was performed in GraphPad Prism via a two-way ANOVA with multiple comparisons between day 0 and 28 time points. No asterisks indicate no significance. *** indicates p<0.001, **** indicates p<0.0001.

**S1 Table. Input of parasite lines into the AreBar.**

**S2 Table. Resistant lines against PMX-selective inhibitors generated in this study.**

**S3 Table. Summary of mutations found by whole genome sequencing.**

**S4 Table. EC_50_ values of resistant and reversion lines against PMX selective and PMIX/X dual inhibitors.**

## References

1. WHO. World malaria report 2025. 2025.

2. White NJ, Chotivanich K. Artemisinin-resistant malaria. Clin Microbiol Rev. 2024;37(4):e0010924. Epub 20241015. doi: 10.1128/cmr.00109-24. PubMed PMID: 39404268; PubMed Central PMCID: PMCPMC11629630.

3. Dhorda M, Kaneko A, Komatsu R, Kc A, Mshamu S, Gesase S, et al. Artemisinin-resistant malaria in Africa demands urgent action. Science. 2024;385(6706):252–4. Epub 20240718. doi: 10.1126/science.adp5137. PubMed PMID: 39024426.

4. Ariey F, Witkowski B, Amaratunga C, Beghain J, Langlois AC, Khim N, et al. A molecular marker of artemisinin-resistant Plasmodium falciparum malaria. Nature. 2014;505(7481):50–5. doi: 10.1038/nature12876. PubMed PMID: 24352242.

5. Wicht KJ, Mok S, Fidock DA. Molecular mechanisms of drug resistance in Plasmodium falciparum malaria. Annu Rev Microbiol. 2020;74:431–54. doi: 10.1146/annurev-micro-020518-115546. PubMed PMID: 32905757; PubMed Central PMCID: PMCPMC8130186.

6. Behrens HM, Schmidt S, Spielmann T. The newly discovered role of endocytosis in artemisinin resistance. Med Res Rev. 2021;41(6):2998–3022. Epub 20210726. doi: 10.1002/med.21848. PubMed PMID: 34309894.

7. Witkowski B, Duru V, Khim N, Ross LS, Saintpierre B, Beghain J, et al. A surrogate marker of piperaquine-resistant Plasmodium falciparum malaria: a phenotype-genotype association study. Lancet Infect Dis. 2017;17(2):174–83. Epub 20161103. doi: 10.1016/S1473-3099(16)30415-7. PubMed PMID: 27818097; PubMed Central PMCID: PMCPMC5266792.

8. Amato R, Lim P, Miotto O, Amaratunga C, Dek D, Pearson RD, et al. Genetic markers associated with dihydroartemisinin-piperaquine failure in Plasmodium falciparum malaria in Cambodia: a genotype-phenotype association study. Lancet Infect Dis. 2017;17(2):164–73. Epub 20161103. doi: 10.1016/S1473-3099(16)30409-1. PubMed PMID: 27818095; PubMed Central PMCID: PMCPMC5564489.

9. van der Pluijm RW, Imwong M, Chau NH, Hoa NT, Thuy-Nhien NT, Thanh NV, et al. Determinants of dihydroartemisinin-piperaquine treatment failure in Plasmodium falciparum malaria in Cambodia, Thailand, and Vietnam: a prospective clinical, pharmacological, and genetic study. Lancet Infect Dis. 2019;19(9):952–61. Epub 20190722. doi: 10.1016/S1473-3099(19)30391-3. PubMed PMID: 31345710; PubMed Central PMCID: PMCPMC6715822.

10. Deni I, Stokes BH, Ward KE, Fairhurst KJ, Pasaje CFA, Yeo T, et al. Mitigating the risk of antimalarial resistance via covalent dual-subunit inhibition of the Plasmodium proteasome. Cell Chem Biol. 2023;30(5):470–85 e6. Epub 20230323. doi: 10.1016/j.chembiol.2023.03.002. PubMed PMID: 36963402; PubMed Central PMCID: PMCPMC10198959.

11. Mohammed R, Asres MS, Gudina EK, Adissu W, Johnstone H, Marrast AC, et al. Efficacy, safety, tolerability, and pharmacokinetics of MMV390048 in acute uncomplicated malaria. Am J Trop Med Hyg. 2023;108(1):81–4. Epub 20221205. doi: 10.4269/ajtmh.22-0567. PubMed PMID: 36509063; PubMed Central PMCID: PMCPMC9833083.

12. Phillips MA, Lotharius J, Marsh K, White J, Dayan A, White KL, et al. A long-duration dihydroorotate dehydrogenase inhibitor (DSM265) for prevention and treatment of malaria. Sci Transl Med. 2015;7(296):296ra111. Epub 2015/07/17. doi: 10.1126/scitranslmed.aaa6645. PubMed PMID: 26180101; PubMed Central PMCID: PMCPMC4539048.

13. Sangana R, Ogutu B, Yeka A, Kusemererwa S, Tinto H, Toure AO, et al. Pharmacokinetics of ganaplacide and lumefantrine in adults, adolescents, and children with Plasmodium falciparum malaria treated with ganaplacide plus lumefantrine solid dispersion formulation: analysis of data from a Multinational Phase 2 Study. J Clin Pharmacol. 2025;65(2):179–89. Epub 20240929. doi: 10.1002/jcph.6138. PubMed PMID: 39344281; PubMed Central PMCID: PMCPMC11771541.

14. Siqueira-Neto JL, Wicht KJ, Chibale K, Burrows JN, Fidock DA, Winzeler EA. Antimalarial drug discovery: progress and approaches. Nat Rev Drug Discov. 2023;22(10):807–26. Epub 20230831. doi: 10.1038/s41573-023-00772-9. PubMed PMID: 37652975; PubMed Central PMCID: PMCPMC10543600.

15. Pino P, Caldelari R, Mukherjee B, Vahokoski J, Klages N, Maco B, et al. A multistage antimalarial targets the plasmepsins IX and X essential for invasion and egress. Science. 2017;358(6362):522–8. Epub 2017/10/28. doi: 10.1126/science.aaf8675. PubMed PMID: 29074775; PubMed Central PMCID: PMCPMC5730047.

16. Nasamu AS, Glushakova S, Russo I, Vaupel B, Oksman A, Kim AS, et al. Plasmepsins IX and X are essential and druggable mediators of malaria parasite egress and invasion. Science. 2017;358(6362):518–22. Epub 2017/10/28. doi: 10.1126/science.aan1478. PubMed PMID: 29074774; PubMed Central PMCID: PMCPMC5928414.

17. Favuzza P, de Lera Ruiz M, Thompson JK, Triglia T, Ngo A, Steel RWJ, et al. Dual plasmepsin-targeting antimalarial agents disrupt multiple stages of the malaria parasite life cycle. Cell Host Microbe. 2020;27(4):642–58 e12. Epub 2020/02/29. doi: 10.1016/j.chom.2020.02.005. PubMed PMID: 32109369; PubMed Central PMCID: PMCPMC7146544.

18. de Lera Ruiz M, Favuzza P, Guo Z, Zhao L, Hu B, Lei Z, et al. The invention of WM382, a highly potent PMIX/X dual inhibitor toward the treatment of malaria. ACS Med Chem Lett. 2022;13(11):1745–54. Epub 20221012. doi: 10.1021/acsmedchemlett.2c00355. PubMed PMID: 36385924; PubMed Central PMCID: PMCPMC9661708.

19. Favuzza P, Palandri J, de Lera Ruiz M, Bailey W, Boyce CW, Danziger A, et al. MK-7602: a potent multi-stage dual-targeting antimalarial. EBioMedicine. 2025;123:106061. Epub 20251206. doi: 10.1016/j.ebiom.2025.106061. PubMed PMID: 41353980.

20. Scally SW, Triglia T, Evelyn C, Seager BA, Pasternak M, Lim PS, et al. PCRCR complex is essential for invasion of human erythrocytes by Plasmodium falciparum. Nat Microbiol. 2022;7(12):2039–53. Epub 20221117. doi: 10.1038/s41564-022-01261-2. PubMed PMID: 36396942; PubMed Central PMCID: PMCPMC9712106.

21. Triglia T, Scally SW, Seager BA, Pasternak M, Dagley LF, Cowman AF. Plasmepsin X activates the PCRCR complex of Plasmodium falciparum by processing PfRh5 for erythrocyte invasion. Nat Commun. 2023;14(1):2219. Epub 20230419. doi: 10.1038/s41467-023-37890-2. PubMed PMID: 37072430; PubMed Central PMCID: PMCPMC10113190.

22. Hodder AN, Christensen J, Scally S, Triglia T, Ngo A, Birkinshaw RW, et al. Basis for drug selectivity of plasmepsin IX and X inhibition in Plasmodium falciparum and vivax. Structure. 2022;30(7):947–61 e6. Epub 20220422. doi: 10.1016/j.str.2022.03.018. PubMed PMID: 35460613.

23. Stanley SE, Carstens RP, Liberti MV, Eertmans W, Vranckx M, Longo D, et al. First-in-human safety and pharmacokinetics of MK-7602, the antimalarial inhibitor of plasmepsins IX/X, in single- and multiple-ascending-dose studies. Antimicrob Agents Chemother. 2026:e0126125. Epub 20260114. doi: 10.1128/aac.01261-25. PubMed PMID: 41532789.

24. Rottmann M, McNamara C, Yeung BK, Lee MC, Zou B, Russell B, et al. Spiroindolones, a potent compound class for the treatment of malaria. Science. 2010;329(5996):1175–80. Epub 2010/09/04. doi: 329/5996/1175 [pii] 10.1126/science.1193225. PubMed PMID: 20813948.

25. Lim MY, LaMonte G, Lee MCS, Reimer C, Tan BH, Corey V, et al. UDP-galactose and acetyl-CoA transporters as Plasmodium multidrug resistance genes. Nat Microbiol. 2016;1:16166. Epub 20160919. doi: 10.1038/nmicrobiol.2016.166. PubMed PMID: 27642791; PubMed Central PMCID: PMCPMC5575994.

26. Carrasquilla M, Drammeh NF, Rawat M, Sanderson T, Zenonos Z, Rayner JC, et al. Barcoding genetically distinct Plasmodium falciparum strains for comparative assessment of fitness and antimalarial drug resistance. mBio. 2022;13(5):e0093722. Epub 20220816. doi: 10.1128/mbio.00937-22. PubMed PMID: 35972144; PubMed Central PMCID: PMCPMC9600763.

27. Qiu D, Pei JV, Rosling JEO, Thathy V, Li D, Xue Y, et al. A G358S mutation in the Plasmodium falciparum Na+ pump PfATP4 confers clinically-relevant resistance to cipargamin. Nat Commun. 2022;13(1):5746. Epub 20220930. doi: 10.1038/s41467-022-33403-9. PubMed PMID: 36180431; PubMed Central PMCID: PMCPMC9525273.

28. Barnes DA, Foote SJ, Galatis D, Kemp DJ, Cowman AF. Selection for high-level chloroquine resistance results in deamplification of the pfmdr1 gene and increased sensitivity to mefloquine in Plasmodium falciparum. EMBO J. 1992;11(8):3067–75.

29. Cowman AF, Galatis D, Thompson JK. Selection for mefloquine resistance in Plasmodium falciparum is linked to amplification of the pfmdr1 gene and cross-resistance to halofantrine and quinine. Proc Natl Acad Sci U S A. 1994;91(3):1143–7. doi: 10.1073/pnas.91.3.1143. PubMed PMID: 8302844; PubMed Central PMCID: PMCPMC521470.

30. Rathod PK, McErlean T, Lee P-C. Variations in frequencies of drug resistance in Plasmodium falciparum. Proc Natl Acad Sci USA. 1997;94:9389-93.

31. Cowell AN, Istvan ES, Lukens AK, Gomez-Lorenzo MG, Vanaerschot M, Sakata-Kato T, et al. Mapping the malaria parasite druggable genome by using in vitro evolution and chemogenomics. Science. 2018;359(6372):191–9. doi: 10.1126/science.aan4472. PubMed PMID: 29326268; PubMed Central PMCID: PMCPMC5925756.

32. Lowe MA, Cardenas A, Valentin JP, Zhu Z, Abendroth J, Castro JL, et al. Discovery and characterization of potent, efficacious, orally available antimalarial plasmepsin X inhibitors and preclinical safety assessment of UCB7362. J Med Chem. 2022;65(20):14121–43. Epub 20221010. doi: 10.1021/acs.jmedchem.2c01336. PubMed PMID: 36216349; PubMed Central PMCID: PMCPMC9620073.

33. LaMonte G, Lim MY, Wree M, Reimer C, Nachon M, Corey V, et al. Mutations in the Plasmodium falciparum Cyclic Amine Resistance Locus (pfcarl) Confer Multidrug Resistance. mBio. 2016;7(4). Epub 20160705. doi: 10.1128/mBio.00696-16. PubMed PMID: 27381290; PubMed Central PMCID: PMCPMC4958248.

34. Murithi JM, Pascal C, Bath J, Boulenc X, Gnadig NF, Pasaje CFA, et al. The antimalarial MMV688533 provides potential for single-dose cures with a high barrier to Plasmodium falciparum parasite resistance. Sci Transl Med. 2021;13(603). doi: 10.1126/scitranslmed.abg6013. PubMed PMID: 34290058; PubMed Central PMCID: PMCPMC8530196.

35. Dhingra SK, Small-Saunders JL, Menard D, Fidock DA. Plasmodium falciparum resistance to piperaquine driven by PfCRT. Lancet Infect Dis. 2019;19(11):1168–9. doi: 10.1016/S1473-3099(19)30543-2. PubMed PMID: 31657776; PubMed Central PMCID: PMCPMC6943240.

36. Rhee SY, Grant PM, Tzou PL, Barrow G, Harrigan PR, Ioannidis JPA, et al. A systematic review of the genetic mechanisms of dolutegravir resistance. J Antimicrob Chemother. 2019;74(11):3135–49. doi: 10.1093/jac/dkz256. PubMed PMID: 31280314; PubMed Central PMCID: PMCPMC6798839.

37. Price RN, Uhlemann AC, Brockman A, McGready R, Ashley E, Phaipun L, et al. Mefloquine resistance in Plasmodium falciparum and increased pfmdr1 gene copy number. Lancet. 2004;364(9432):438–47. doi: 10.1016/S0140-6736(04)16767-6. PubMed PMID: 15288742; PubMed Central PMCID: PMCPMC4337987.

38. Wensing AM, Calvez V, Ceccherini-Silberstein F, Charpentier C, Gunthard HF, Paredes R, et al. 2022 update of the drug resistance mutations in HIV-1. Top Antivir Med. 2022;30(4):559–74. PubMed PMID: 36375130; PubMed Central PMCID: PMCPMC9681141.

39. Stokes BH, Dhingra SK, Rubiano K, Mok S, Straimer J, Gnadig NF, et al. Plasmodium falciparum K13 mutations in Africa and Asia impact artemisinin resistance and parasite fitness. Elife. 2021;10. Epub 20210719. doi: 10.7554/eLife.66277. PubMed PMID: 34279219; PubMed Central PMCID: PMCPMC8321553.

40. Trager W, Jensen JB. Human malaria parasites in continuous culture. Science. 1976;193:673–5.

41. Gamo FJ, Sanz LM, Vidal J, de Cozar C, Alvarez E, Lavandera JL, et al. Thousands of chemical starting points for antimalarial lead identification. Nature. 2010;465(7296):305–10. Epub 2010/05/21. doi: nature09107 [pii] 10.1038/nature09107. PubMed PMID: 20485427.

42. Mukherjee S, Nguyen S, Sharma E, Goldberg DE. Maturation and substrate processing topography of the Plasmodium falciparum invasion/egress protease plasmepsin X. Nat Commun. 2022;13(1):4537. Epub 20220804. doi: 10.1038/s41467-022-32271-7. PubMed PMID: 35927261; PubMed Central PMCID: PMCPMC9352755.

43. Jain NK, Barkowski-Clark S, Altman R, Johnson K, Sun F, Zmuda J, et al. A high density CHO-S transient transfection system: Comparison of ExpiCHO and Expi293. Protein Expr Purif. 2017;134:38–46. Epub 2017/03/28. doi: 10.1016/j.pep.2017.03.018. PubMed PMID: 28342833.

44. Scheinin I, Sie D, Bengtsson H, van de Wiel MA, Olshen AB, van Thuijl HF, et al. DNA copy number analysis of fresh and formalin-fixed specimens by shallow whole-genome sequencing with identification and exclusion of problematic regions in the genome assembly. Genome Res. 2014;24(12):2022–32. Epub 20140918. doi: 10.1101/gr.175141.114. PubMed PMID: 25236618; PubMed Central PMCID: PMCPMC4248318.

45. Cameron DL, Baber J, Shale C, Valle-Inclan JE, Besselink N, van Hoeck A, et al. GRIDSS2: comprehensive characterisation of somatic structural variation using single breakend variants and structural variant phasing. Genome Biol. 2021;22(1):202. Epub 20210712. doi: 10.1186/s13059-021-02423-x. PubMed PMID: 34253237; PubMed Central PMCID: PMCPMC8274009.

46. Cameron DL, Schroder J, Penington JS, Do H, Molania R, Dobrovic A, et al. GRIDSS: sensitive and specific genomic rearrangement detection using positional de Bruijn graph assembly. Genome Res. 2017;27(12):2050–60. Epub 20171102. doi: 10.1101/gr.222109.117. PubMed PMID: 29097403; PubMed Central PMCID: PMCPMC5741059.

47. Pockrandt C, Alzamel M, Iliopoulos CS, Reinert K. GenMap: ultra-fast computation of genome mappability. Bioinformatics. 2020;36(12):3687–92. doi: 10.1093/bioinformatics/btaa222. PubMed PMID: 32246826; PubMed Central PMCID: PMCPMC7320602.

48. Miles A, Iqbal Z, Vauterin P, Pearson R, Campino S, Theron M, et al. Indels, structural variation, and recombination drive genomic diversity in Plasmodium falciparum. Genome Res. 2016;26(9):1288–99. Epub 20160816. doi: 10.1101/gr.203711.115. PubMed PMID: 27531718; PubMed Central PMCID: PMCPMC5052046

